# Methylome and transcriptome mapping reveal miniscule DNA methyltransferase regulons in *Salmonella enterica* serovar Typhimurium

**DOI:** 10.64898/2026.01.27.702048

**Authors:** Anna S. Ershova, Charlie Howard, Karsten Hokamp, Andrew D.S. Cameron, Carsten Kröger

## Abstract

DNA methylation is a regulator of bacterial gene expression and adaptation, influencing traits such as virulence and antimicrobial resistance. The dynamic nature of DNA methylation enables rapid responses to changing environments and is a source of heterogeneity in bacterial populations. However, condition-dependent DNA methylation and consequences for transcriptional output remain poorly understood. We applied Oxford Nanopore sequencing to profile DNA methylation during exponential growth and late stationary phase of *Salmonella enterica* serovar Typhimurium and integrated these data with transcriptomic analyses. We found that each DNA methyltransferase (MTases) exhibits a distinct activity pattern across growth stages, which could not be explained by transcriptional levels of the corresponding enzymes. As predicted, DNA methylation patterns determined by regulatory MTases were dynamic across growth conditions whereas methylation patterns of MTases belonging to R-M systems were comparatively stable. We identified growth stage–specific methylation patterns for all studied MTases and correlations between methylation states and gene expression patterns. Together, these findings chart DNA methylation networks in the epigenetic regulation of bacterial physiology.

**Author summary:** DNA methylation in bacteria is best known for its role protecting DNA from endonucleases, such as restriction–modification, and coordinating chromosome replication and mutation repair, yet DNA methylation also regulates gene expression and cell physiology. Previous studies primarily examined bacterial DNA methylation at single time points or in limited genomic regions, providing only a partial view of its biological significance. In this study, we used Oxford Nanopore sequencing to compare DNA methylation patterns in *Salmonella enterica* during exponential growth and late stationary phase then integrated these data with corresponding gene expression profiles. We identified numerous methylation target motifs, all of which demonstrated constitutively methylated or unmethylated regions. This systems-level analysis clarifies the role of DNA methylation in bacterial adaptation across growth stages and demonstrates the utility of Oxford Nanopore sequencing for genome-wide methylation profiling.

## Introduction

Bacterial DNA methylation occurs in several forms, including N6-methyladenine (6mA), 5-methylcytosine (5mC), and N4-methylcytosine (4mC) and is carried out by DNA methyltransferases (MTases) [1,2]. MTases occur either as solitary enzymes or as part of defence systems. Solitary MTases often function in regulating gene expression or other cellular processes [3–5]. Many MTases are components of antiviral defence systems, most notably Restriction–Modification (R–M) systems [6]. R-M systems are classified into Types I, II, and III according to their enzyme architecture, DNA-recognition site structure, and required cofactors [7]. Defence-associated MTases also can contribute to epigenetic regulation, linking their protective function to broader control of bacterial physiology [8].

*Salmonella enterica* encodes several R-M systems and three regulatory MTases: m6A MTases Dam methylating GATC and YhdJ methylating ATGCAT, and m5C MTase Dcm, methylating CCWGG [7]. The regulatory role of GATC methylation has been well studied in *S. enterica*, where Dam-dependent regulation of the *opvAB* operon controls O-antigen chain length, affecting phage resistance and immune evasion [9]. Similarly, the *gtr* operon encodes glucosyltransferases responsible for O-antigen modification, promoting phage resistance [10]. Dam also regulates the *std* fimbrial operon, which influences adhesion and intestinal colonization [11]. Changes in GATC methylation states in promoter regions can generate subpopulations with distinct expression profiles, contributing to phenotypic heterogeneity [12]. Dcm methylation is growth-phase dependent; many CCWGG sites are more frequently methylated in stationary phase compared to exponential phase in *E. coli* [13]. Deletion of *dcm* affects gene expression, particularly in stationary phase, but presence of a CCWGG motif in upstream regions does not faithfully predict expression changes [13]. Compared to Dam and Dcm, the function of YhdJ remains unclear [14,15].

DNA methylation states can be detected by DNA sequencing platforms that sequence native DNA molecules. In bacteria, PacBio SMRT sequencing reliably identifies 6mA and 4mC but not 5mC methylation [16]. Because detection of 5mC typically requires bisulfite sequencing, comprehensive bacterial 5mC datasets are relatively rare [13]. Oxford Nanopore sequencing has the potential to detect all methylated bases, enabling in-depth analysis of bacterial methylomes [17]; however, it remains unclear how Oxford Nanopore–derived methylation data benchmarks against PacBio data at the resolution of individual genomic positions.

Methylation states are dynamic, yet most bacterial methylome studies have focused on a single growth phase. Though informative, this static view provides limited insights into epigenetic regulation and dynamics of DNA methylation. Comparative analyses of methylation states in *E. coli* [13,18,19], *Y. pestis* [20], and *S. enterica* [15] examined how methylation states can respond to changing environmental conditions. However, integrating SMRT-based methylome and RNA-seq based transcriptome analyses suggested that methylation states in *S. enterica* are only weakly associated with transcriptional outputs [15].

In this work, we used Oxford Nanopore sequencing to profile the methylome and methylation dynamics of *S. enterica* 4/74 at mid-exponential (MEP) and late stationary phase (LSP). We hypothesized each MTase has a distinct regulon, which we addressed by cataloguing the methylation states and transcription levels of all putative methylation targets in the two growth states.

### 1. Methyltransferase and Restriction–Modification System Gene Expression across Growth Phases

In this work, we define a methylation motif as the consensus nucleotide sequence recognized and methylated by a specific DNA methyltransferase (e.g., GATC for Dam MTase). By contrast, a methylation site is an individual genomic occurrence of such a motif, for which the methylation status was assessed. According to the REBASE database [7], *S. enterica* 4/74 encodes eleven MTases: three MTases of R-M systems and eight solitary MTases (**Table 1**). To evaluate the expression of MTase genes, we analysed published RNA-seq data for *S. enterica* strain 4/74 obtained from the mid-exponential (MEP) and late stationary (LSP) growth phases (GSE49829, **Supplementary Table 1**) [21].

**Table 1.**
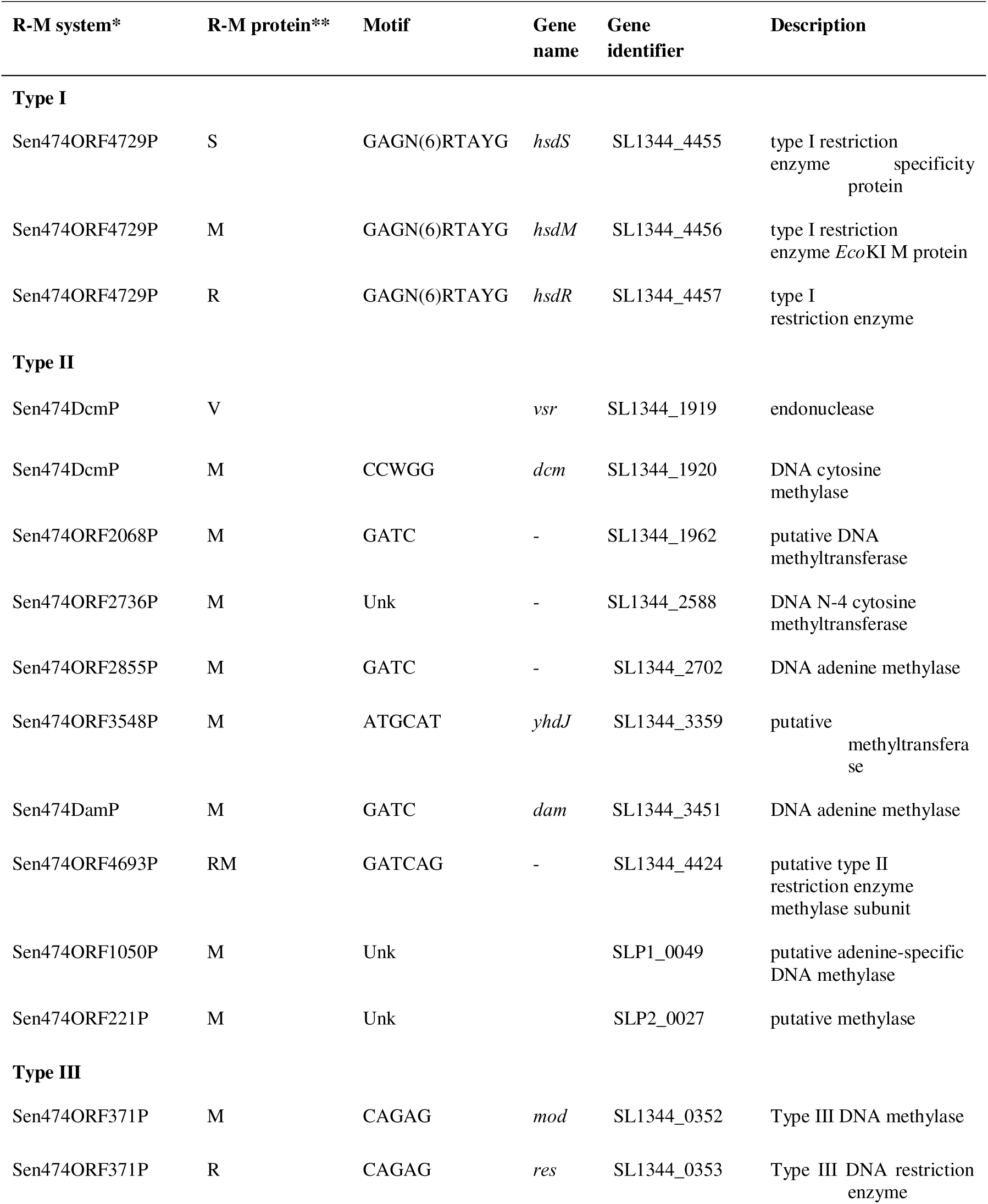
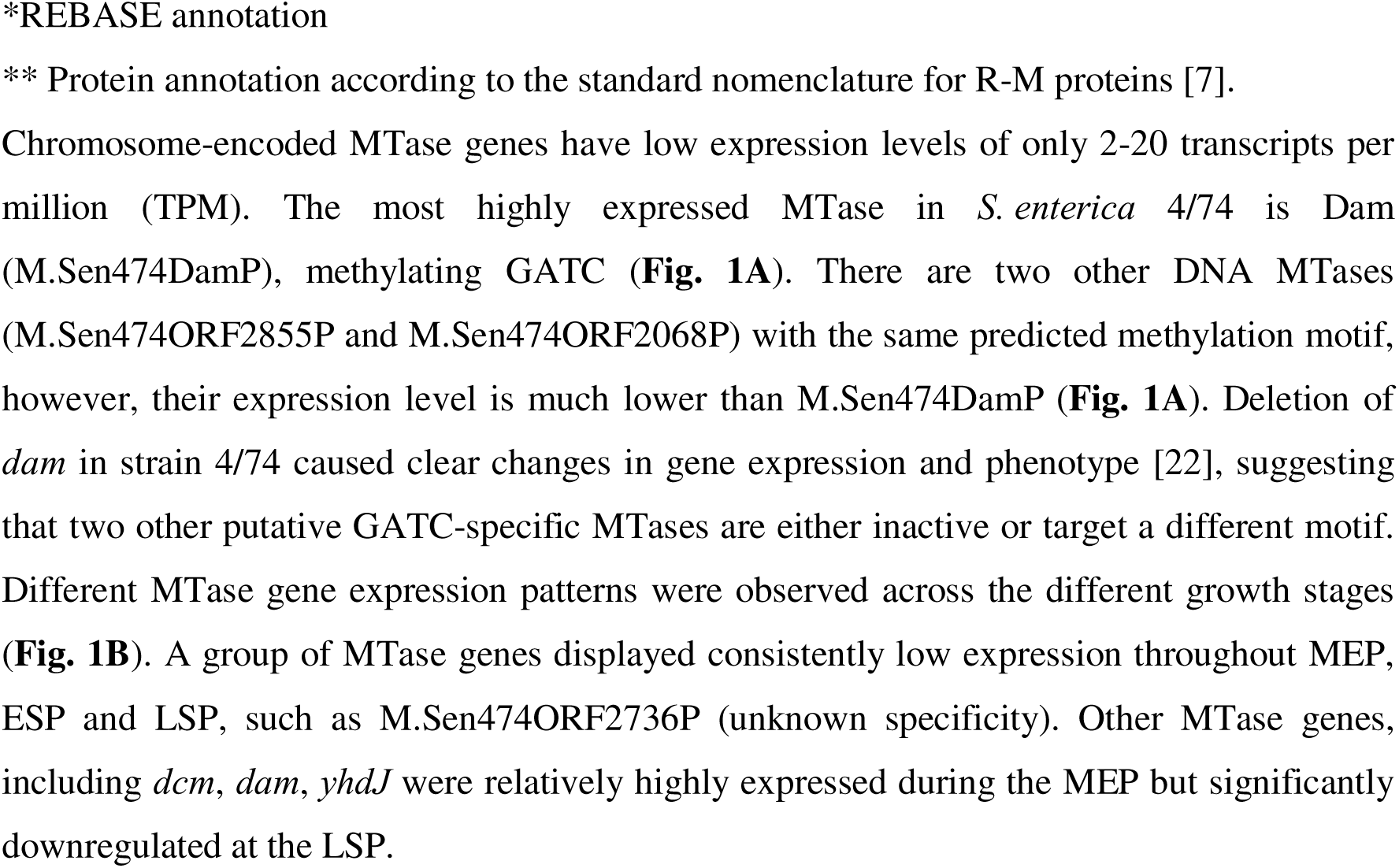
R-M system genes encoded in the *S. enterica* 4/74 genome.

| R-M system* | R-M protein** | Motif | Gene name | Gene identifier | Description |
| --- | --- | --- | --- | --- | --- |
| **Type I** |  |  |  |  |  |
| Sen474ORF4729P | S | GAGN(6)RTAYG | *hsdS* | SL1344_4455 | type I restriction enzyme specificity protein |
| Sen474ORF4729P | M | GAGN(6)RTAYG | *hsdM* | SL1344_4456 | type I restriction enzyme *EcoKI* M protein |
| Sen474ORF4729P | R | GAGN(6)RTAYG | *hsdR* | SL1344_4457 | type I restriction enzyme |
| **Type II** |  |  |  |  |  |
| Sen474DcmP | V |  | *vsr* | SL1344_1919 | endonuclease |
| Sen474DcmP | M | CCWGG | *dcm* | SL1344_1920 | DNA cytosine methylase |
| Sen474ORF2068P | M | GATC | - | SL1344_1962 | putative DNA methyltransferase |
| Sen474ORF2736P | M | Unk | - | SL1344_2588 | DNA N-4 cytosine methyltransferase |
| Sen474ORF2855P | M | GATC | - | SL1344_2702 | DNA adenine methylase |
| Sen474ORF3548P | M | ATGCAT | *yhdJ* | SL1344_3359 | putative methyltransferase |
| Sen474DamP | M | GATC | *dam* | SL1344_3451 | DNA adenine methylase |
| Sen474ORF4693P | RM | GATCAG | - | SL1344_4424 | putative type II restriction enzyme methylase subunit |
| Sen474ORF1050P | M | Unk |  | SLP1_0049 | putative adenine-specific DNA methylase |
| Sen474ORF221P | M | Unk |  | SLP2_0027 | putative methylase |
| **Type III** |  |  |  |  |  |
| Sen474ORF371P | M | CAGAG | *mod* | SL1344_0352 | Type III DNA methylase |
| Sen474ORF371P | R | CAGAG | *res* | SL1344_0353 | Type III DNA restriction enzyme |

\*REBASE annotation
\*\* Protein annotation according to the standard nomenclature for R-M proteins [7].

### 2. Identification of *S. enterica* 4/74 Methylation Motifs

Before conducting a genome-wide assessment of methylation sites and methylation states, we first assessed the sensitivity of methylation detection using Oxford Nanopore DNA sequencing to establish an appropriate calling threshold. To this end, we performed an *in vitro* genomic DNA methylation experiment using M.EcoRI, which specifically methylates GAATTC sites and is not encoded in *S. enterica* 4/74; and then determined the sequencing depth required to detect methylated sites (**Supplementary File 1**). A methylation detection threshold of 35% provided optimal discrimination between methylated and unmethylated sites (**Supplementary File 1**). Methyl group detection improved markedly between 5× and 30× fold sequencing coverage but increased gradually from 30× to 90×. In subsequent analyses, we applied the 35% threshold to identify non-methylated sites and calculated a weighted average methylation level for each site across samples to account for differences in sequencing depth.

We applied these established parameters to detect methylation of isolated DNA of wild-type *S. enterica* 4/74 grown in L-broth to MEP and LSP. All six previously reported methylation motifs for both m6A and m5C in *S. enterica* 4/74 were identified (**Table 1; Supplementary Tables 2, 3 and 4**). These motifs were consistently detected across all samples, except for the ATGCAT motif, which was observed only in one MEP sample (32× Samtools sequencing depth, MEP-1) and one LSP (124× Samtools sequencing depth, LSP-1) sample but not in the lower-coverage MEP (16×, MEP-2) and LSP (24×, LSP-2) samples. In addition, this motif is only partially methylated across the genome (**Supplementary Table 4**). These results suggest that sequencing depth influences the ability to detect low-abundance methylation motifs. Nevertheless, even at a sequencing depth of 16×, five of the six known methylated motifs were successfully identified in the MEP-1 sample, demonstrating the robustness of Oxford Nanopore sequencing for methylation detection.

We compared the methylation levels of individual sites in our samples with publicly available PacBio data for *S. enterica* serovar Typhimurium LT2 (coverage ∼230×) obtained from the REBASE database [7], noting that the growth phase for the PacBio sample is not specified. Overall, the PacBio data showed concordance with our Oxford Nanopore results, supporting the reliability of methylation detection in this study (**Supplementary Table 4**).

Additionally, we identified two novel methylation motifs: GATCGNTAT (m6A) and GATCNTC (m4C) (**Supplementary Tables 2 and 3**). *S. enterica* 4/74 encodes two MTase genes with unknown specificities, as well as two annotated GATC-specific MTases (**Table 1**). Further experimental work is required to validate the specificities of these enzymes and confirm whether the two newly identified motifs represent distinct methylation patterns. Both motifs contain the GATC recognition site of Dam methyltransferase (M.Sen474DamP), suggesting that these may represent miscalls arising from GATC methylation. This interpretation is supported by the detection of other diverse GATC-containing motifs in individual samples (**Supplementary Tabl**e **3**). This potential motif overprediction should be considered when analysing uncharacterised bacterial strains. In this study, these additional motifs were excluded from downstream analysis.

#### 2.1. GATC Methylation dynamics across growth phases

In *S. enterica* 4/74, three MTases can methylate adenine of GATC sites according to the REBASE database prediction (**Table 1**). The *dam* gene is the most highly expressed MTase at the MEP (**Fig. 1A**), which likely accounts for the constitutive methylation of almost all chromosomal and plasmidic GATC sites (**Fig. 2A**, **Fig. 2B and C, Supplementary Tables 4 and 5**). The results agree with GATC methylation data for *E. coli,* where most GATC sites are methylated at both MEP and LSP [23]. We found that 43 GATC sites on the chromosome and eight sites on plasmids are never methylated in either growth stage (UM, **Fig. 2C**).

**Fig 1.**
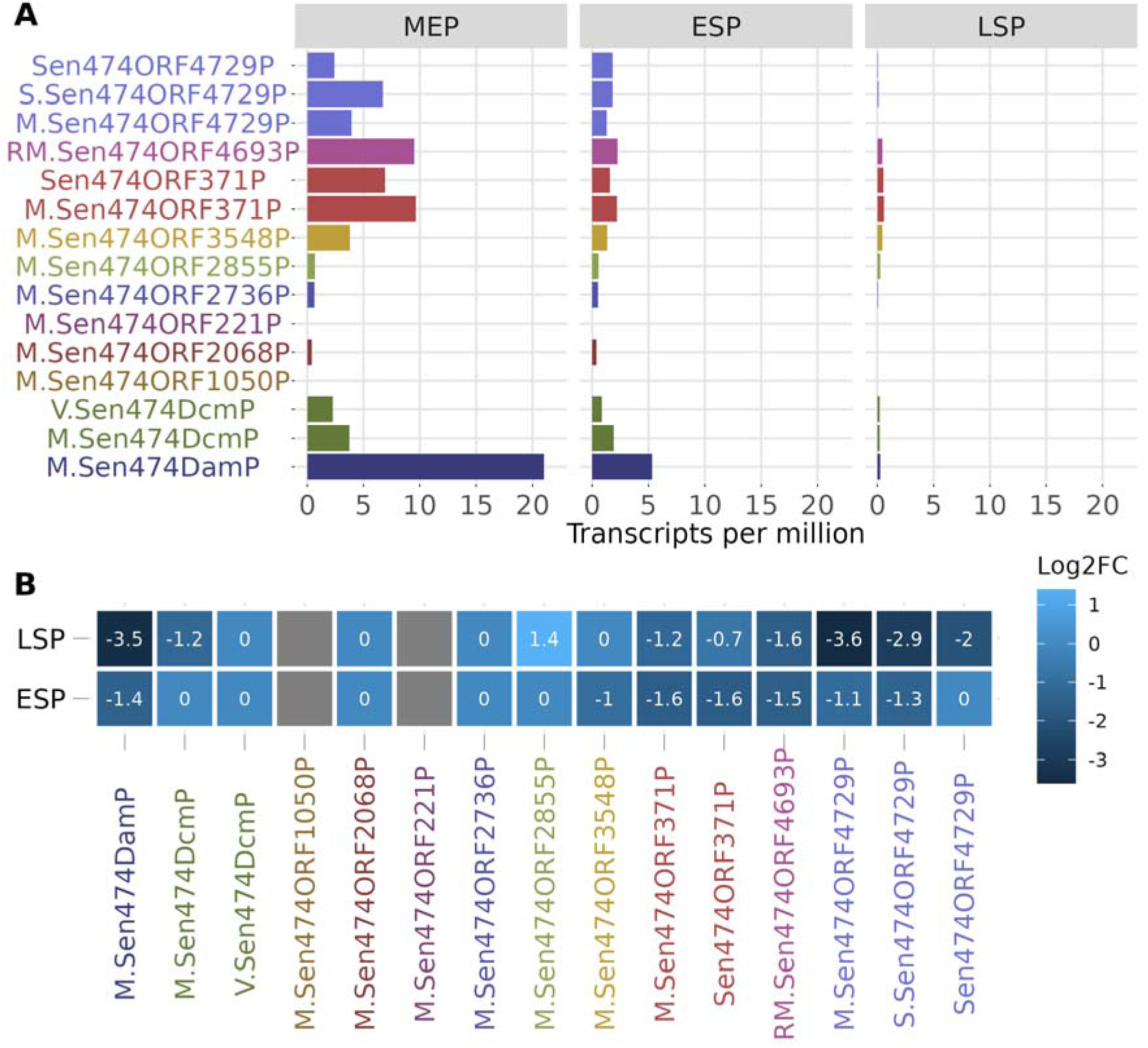
Gene expression of R-M genes at different growth phases (two biological replicas for each phase). MEP-mid exponential phase, ESP-early stationary phase, LSP-late stationary phase. **A**. Transcripts per million (TPM) **B.** DESeq2 results, MEP was used as a reference. Components of the same R-M system are coloured in the same colour. Log_2_FC are shown only if pvalue adjusted <0.01. The intensity of the blue colour shows log_2_FC, grey shows lacking gene expression data.

**Fig 2.**
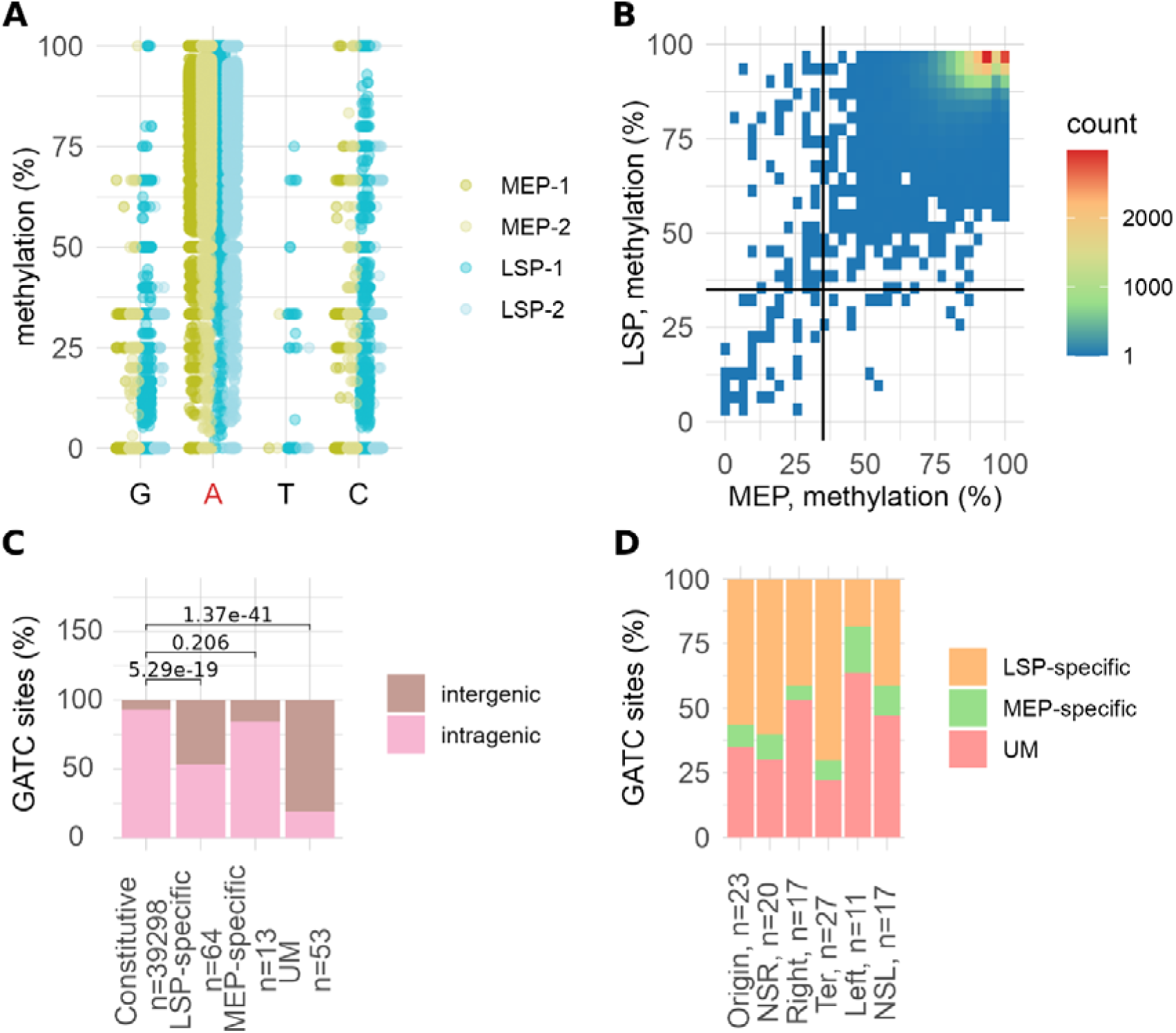
GATC methylation in *S. enterica* 4/74 at different growth phases. **A.** Methylation percentage of bases in the GATC sites across the genome. The A is predominantly methylated (highlighted in red). Two samples of the Mid-exponential phase (MEP-1 and MEP-2) and two samples of the Late Stationary Phase (LSP-1 and LSP-2) are shown. **B.** Methylation dynamics of GATC sites (n=39,428) in a genome across growth phases. 35% methylation thresholds are shown by solid lines. **C.** Distribution of GATC sites by methylation status across intragenic and intergenic regions. LSP-specific and unmethylated sites (UM) are significantly enriched in intergenic regions, Fisher’s exact test, pvalue 5x10^-19^ and 1.37x10^-41^, respectively. **D.** Differentially methylated GATC sites grouped by chromosome macrodomains (NSR = non-structured right, NSL = non-structured left, Ter = terminus). Constitutively methylated sites are omitted. Methylation % corresponds to the percentage of reads, where this position is methylated divided by the total number of reads in this position.

Eighty-one GATC sites on the chromosome were differentially methylated between the two growth states (**Fig. 2C**). Of these, 68 sites were methylated in LSP but not in MEP (LSP-specific), while 13 sites were methylated in MEP but not in LSP (MEP-specific, **Fig. 2C**). On the plasmids, three GATC sites were methylated at MEP and four in LSP (**Supplementary Table 5**). Unmethylated and differentially methylated sites were localised more frequently in intergenic regions than expected by chance, suggesting potential for regulation (**Fig. 2C**). The distribution of GATC sites, including differentially methylated sites, was roughly equal across chromosomal macrodomains (6499±360, **Supplementary Table 6**), suggesting that macrodomain condensation states don’t impact global methylation states (**Fig. 4D**).

#### 2.2. ATGCAT Methylation dynamics across growth phases

The ATGCAT sequence is methylated by the m6A MTase YhdJ (M.Sen474ORF3548P, **Table 1**). As this MTase is encoded by an orphan gene and not part of an R-M system, it may play a regulatory role, however its function is unclear [14,15]. The ATGCAT motif is methylated at the second adenine (**Fig. 3A**) and unlike Dam and Dcm sites, more YhdJ sites are methylated at MEP than at LSP (**Fig. 3B**). Only 61.2% of sites were methylated at least in one growth stage (**Supplementary Table 7**) which is consistent with published and PacBio REBASE data (**Fig. 3B & C, Supplementary Table 4**) [14]. To detect specific sequence motifs or contextual features that may contribute to site-specific methylation, we analysed a five bp window up and downstream of motif sequence contexts surrounding the most highly methylated sites (>50% of methylation at both growth stages) but did not identify obvious patterns (**Supplementary File 2**). Unmethylated ATGCAT sites in both conditions are enriched in intergenic regions, again suggesting a potential role for epigenetic regulation (**Fig. 3C**). Constitutively methylated, differentially methylated, and unmethylated ATGCAT sites are evenly distributed across the chromosomal macrodomains, indicating that no macrodomain shows preferential methylation for this motif. (**Fig. 3D, Supplementary Table 6**).

**Fig 3.**
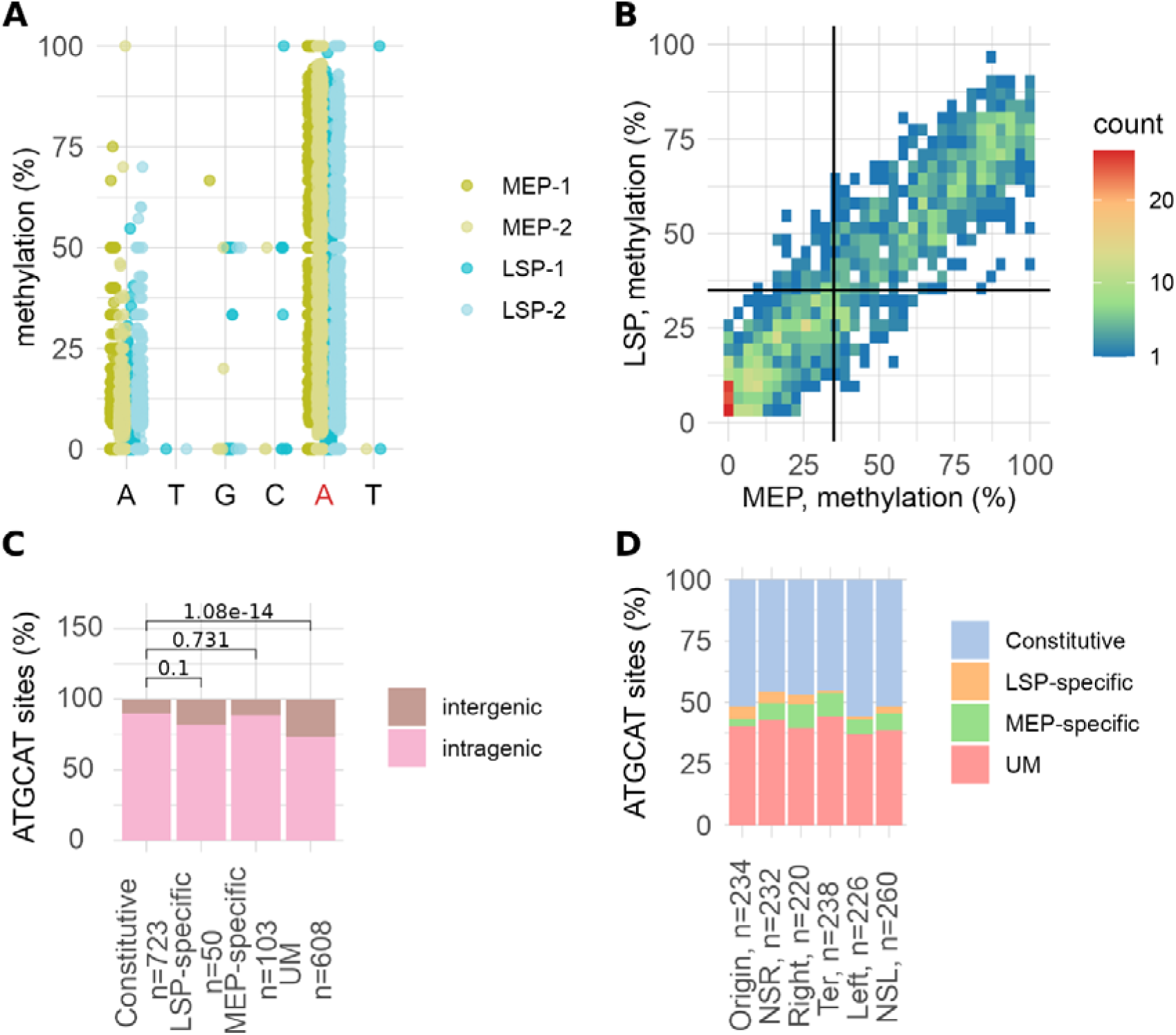
ATGC<u>A</u>T methylation in *S. enterica* 4/74 at different growth phases. **A.** Methylation percentage of bases in the ATGCAT sites across the genome. The second A is methylated (shown in red). Two samples of the Mid-exponential phase (MEP-1 and MEP-2) and two samples of the Late Stationary Phase (LSP-1 and LSP-2) are shown. **B.** Methylation dynamics of ATGCAT sites (n=1,484) in a genome across growth phases. 35% methylation thresholds are shown by solid lines. **C.** Distribution of ATGCAT sites across intragenic regions and intergenic regions grouped by methylation status. Unmethylated sites (UM) are significantly enriched in intergenic regions, chi-square test, pvalue 1x10^-14^ **D.** Distribution of ATGCAT sites across methylation status grouped by chromosome macrodomain. Only chromosomal sites are shown. Methylation % corresponds to the percents of reads, where this position is methylated to the total number of reads in this position.

#### 2.3. CCWGG Methylation dynamics across growth phases

The CCWGG motif is recognised and methylated by Dcm MTase. Here, the second C is predominantly methylated, however, the first C is also identified as frequently methylated (**Fig. 4A**). This effect of a methylated base on neighbouring bases has also been identified previously [24]. The level of CCWGG site methylation is much higher at LSP than at MEP (**Fig. 4B, Supplementary Table 8**) suggesting condition-specific roles for Dcm-mediated methylation. In fact, none of the sites were MEP-specifically methylated as differentially methylated sites were only observed in LSP (**Fig. 4C**). Unmethylated sites are significantly enriched in intergenic regions similar to GATC and ATGCAT motifs (**Fig. 4C**). Finally, the distribution of methylated/unmethylated CCWGG sites was equal across macrodomains (**Fig. 6D**).

**Fig 4.**
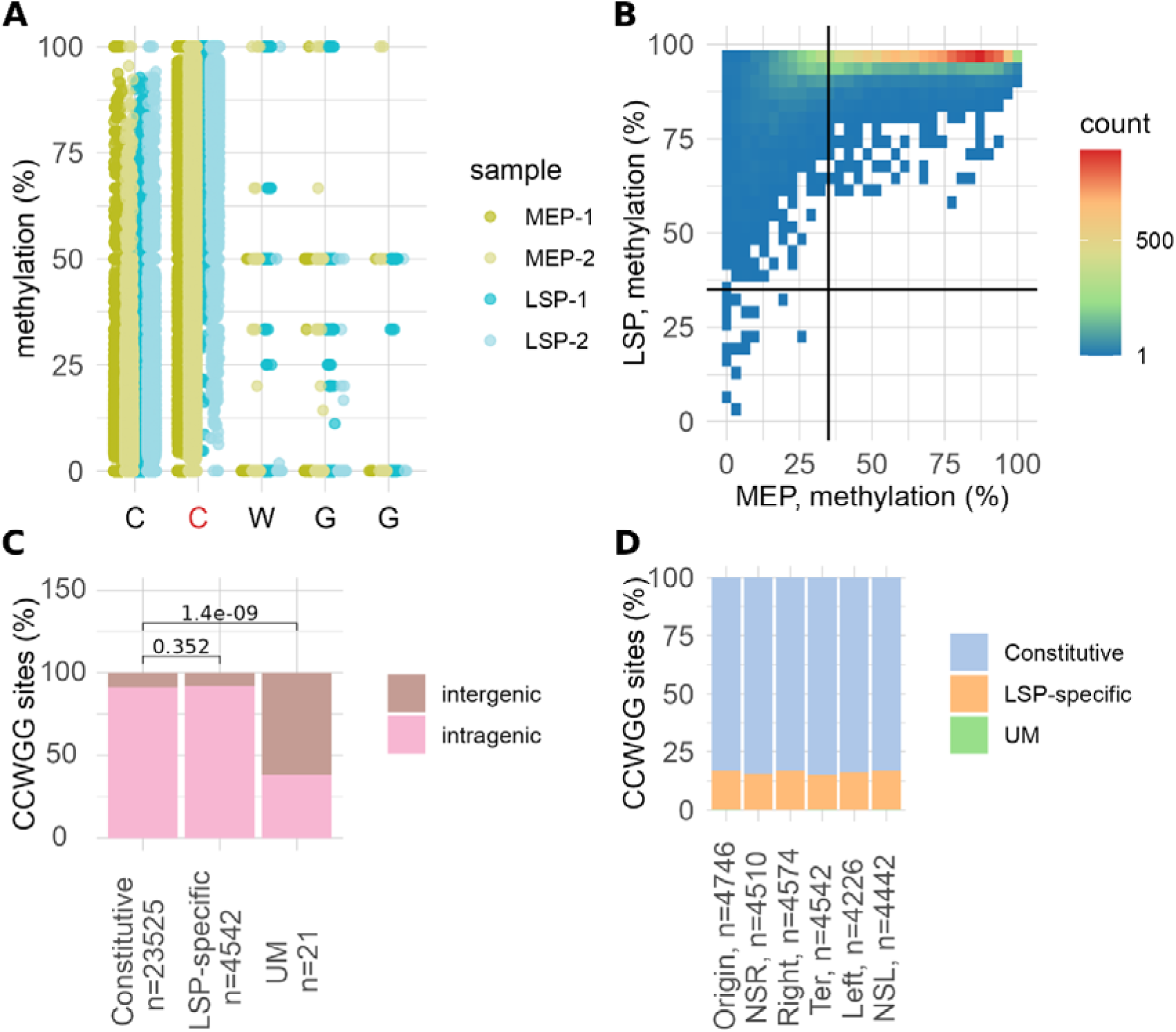
C<u>C</u>WGG methylation in *S. enterica* 4/74 at different growth phases. **A.** Methylation percentage of bases in the CCWGG sites across the genome. The second C is methylated (shown in red). Two samples of the Mid-exponential phase (MEP-1 and MEP-2) and two samples of the Late Stationary Phase (LSP-1 and LSP-2) are shown. **B.** Methylation dynamics of CCWGG sites (n=28,088) in a genome across growth phases. 35% methylation thresholds are shown by solid lines. **C.** Distribution of CCWGG sites across intragenic regions and intergenic regions grouped by methylation status. Unmethylated sites (UM) are significantly enriched in intergenic regions, chi-square test, pvalue =1.4x10^-9^ **D**. Distribution of CCWGG sites across methylation status grouped by chromosome macrodomain. Only chromosomal sites are shown; constitutively methylated sites are omitted. Methylation % corresponds to the percents of reads, where this position is methylated to the total number of reads in this position.

#### 2.4. R-M system MTase methylation

*S. enterica* 4/74 possesses one Type I, one type IIG, and one Type III R-M system. All MTases belonging to R-M systems show a similar methylation pattern, where a majority of sites is constitutively methylated (**Supplementary File 2**). These results correspond to previous data [2] and independently prove the determined methylation threshold of 35%. A relatively low number of sites demonstrate growth-stage dependent methylation pattern.

### 3. Association between differentially methylated sites and differentially expressed genes

We identified condition-specific methylation sites for all three regulatory MTases, including Dam, Dcm, and YhdJ. LSP-specific GATC sites were frequently located in intergenic regions (Fig. 2C), whereas condition-specific methylated ATGCAT (**Fig. 3C**) and CCWGG (**Fig. 4C**) sites were more often found within intragenic regions.

To examine the potential transcriptional impact of condition-dependent methylation, we identified genes whose expression could be affected by differentially methylated sites. For intragenic sites, we selected the genes containing these sites, whereas for intergenic sites, we assigned nearby genes whose expression might be influenced by methylation of their upstream regions. We then compared gene expression changes between LSP and MEP across three categories: (i) genes with condition-specific methylated sites in their upstream regions, (ii) genes with condition-specific methylated sites in intragenic regions, and (iii) all remaining genes, for each of the three regulatory motifs.

We identified 64 LSP-specific GATC sites, including 34 sites in intragenic regions and 30 sites in intergenic regions. LSP-specific GATC sites are often localised in the upstream region 0-250 bp (chi-square p-value < 2.2e-16), where they might be involved in modulation of gene expression (Fig 5B). Among the 13 GATC sites specifically unmethylated in MEP, only two sites occur in intergenic regions, both positioned upstream of the *cdd* gene. We found a significant association between the presence of growth-stage-specific GATC sites in upstream regions and upregulation of the corresponding genes (Mann–Whitney–Wilcoxon test, p = 0.00078). No significant expression changes were observed for genes containing growth-stage-specific methylated GATC sites within intragenic regions (**Fig. 5D**).

**Fig 5.**
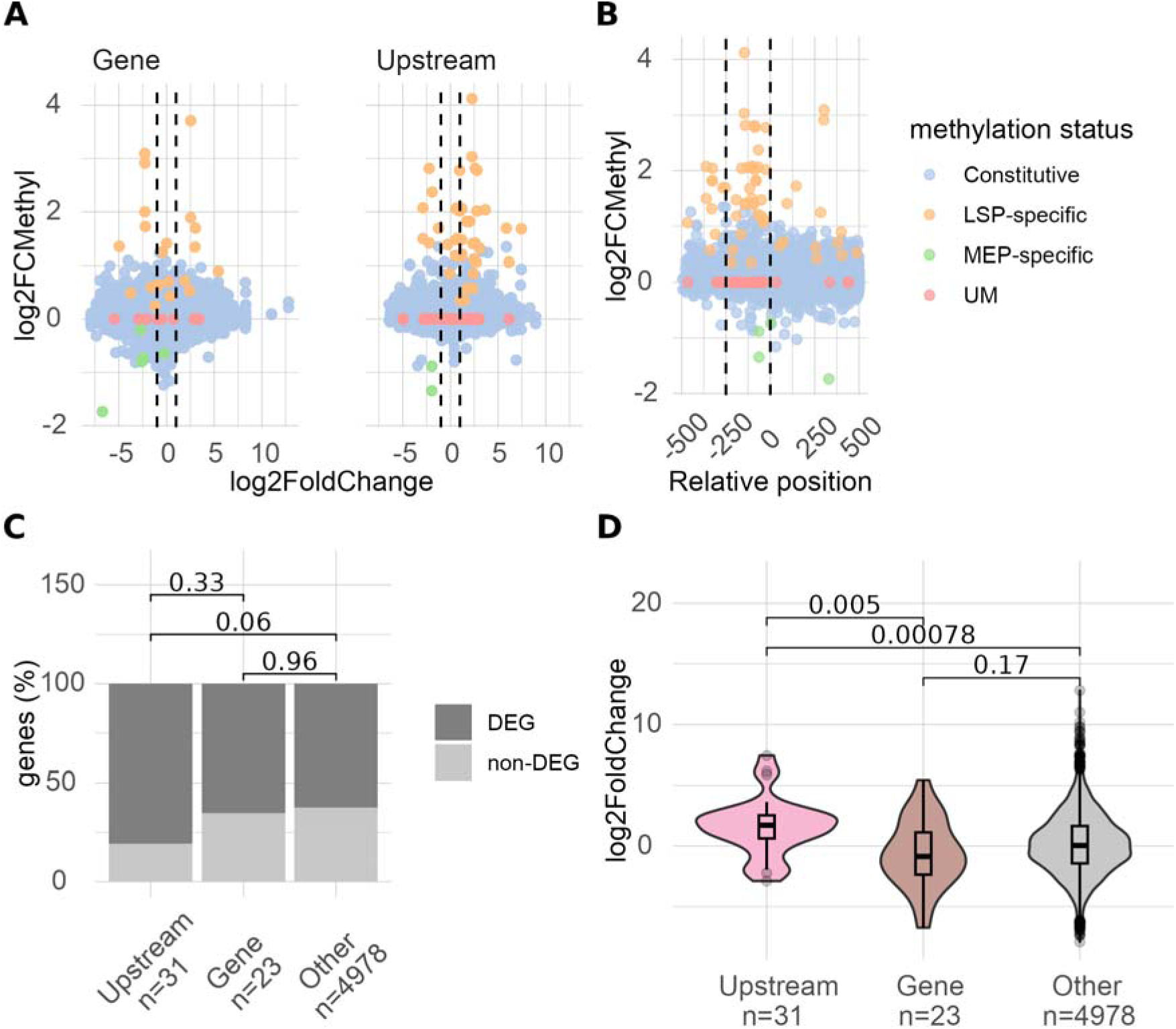
Associations between DNA methylation and gene expression changes between MEP and LSP phases for the GATC motif. **A.** Comparison of changes in gene expression (log□ fold change) and changes in DNA methylation (log_2_FCMethyl) between LSP and MEP for sites localised in intragenic regions (left panel) and upstream regions (right panel). The thresholds for differentially expressed genes are shown as dashed lines (log_2_ fold change is -1 for downregulated and +1 for upregulated genes, padj <0.05) **B.** Relative position of sites (+1 corresponds to the annotated start codon) and changes in DNA methylation between growth phases. Differentially methylated sites are mainly localised in the functionally important upstream area (0-250 bp), chi-square p-value < 2.2e-16. **C.** Qualitative comparison of differentially expressed genes (DEGs) and non-DEGs among groups defined by the presence of differentially methylated sites in upstream regions, intragenic regions, or neither. Genes with a log_2_ fold change ≥ 1 or ≤ –1 and an adjusted p < 0.05 were classified as DEGs; all others were considered non-DEGs. Statistical significance was assessed using a chi-square test. **D.** Quantitative analysis of log_2_ fold change distributions in gene expression for the same groups. Statistical significance was assessed using the Mann–Whitney–Wilcoxon test.

Several LSP-specific sites identified in our dataset overlap with previously characterised regulatory loci, supporting their role in epigenetic control. For example, the LSP-specific GATC sites located upstream of *STM3083* (*STM3726* in *S. enterica* 14028) and *dgoR* correspond to undermethylated sites shown to mediate epigenetic regulation of these genes [12]. Similarly, the GATC site within the *intA* promoter, known to function as an epigenetic switch controlling *intA* expression and prophage ST64B reintegration [25] was also identified as LSP-specific in our dataset (see **Supplementary Table 5**). These examples confirm that a subset of the LSP-specific methylation sites detected here are functionally relevant and likely contribute to epigenetic regulation during the transition from exponential to stationary phase. Future studies may test whether our newly identified intergenic LSP-specific methylation sites impact gene regulation of their downstream genes.

The CCWGG methylation pattern varied significantly between MEP and LSP, with ∼16% of sites (4,884 of 28,088) methylated only at LSP (**Fig. 6, Supplementary Table 8**). No enrichment with LSP- or MEP-specific CCWGG sites in the upstream regions (**Fig. 10 A**, **B**) nor any effect of upstream CCWGG methylation on gene expression was observed (**Fig. 10 C**, **D**). To our knowledge, only one example of condition-specific CCWGG methylation changes has been reported: LSP-specific methylation of the CCWGG site within the *E. coli lexA* repressor promoter [26], which was associated with *lexA* downregulation. However, experiments did not show that methylation directly affected *lexA* expression [26]. In our dataset, we identified this site (positions 4480528–4480533) as LSP-specific and confirmed that *lexA* (SL1344_4174) is downregulated at LSP (see **Supplementary Table 8**).

**Fig 6.**
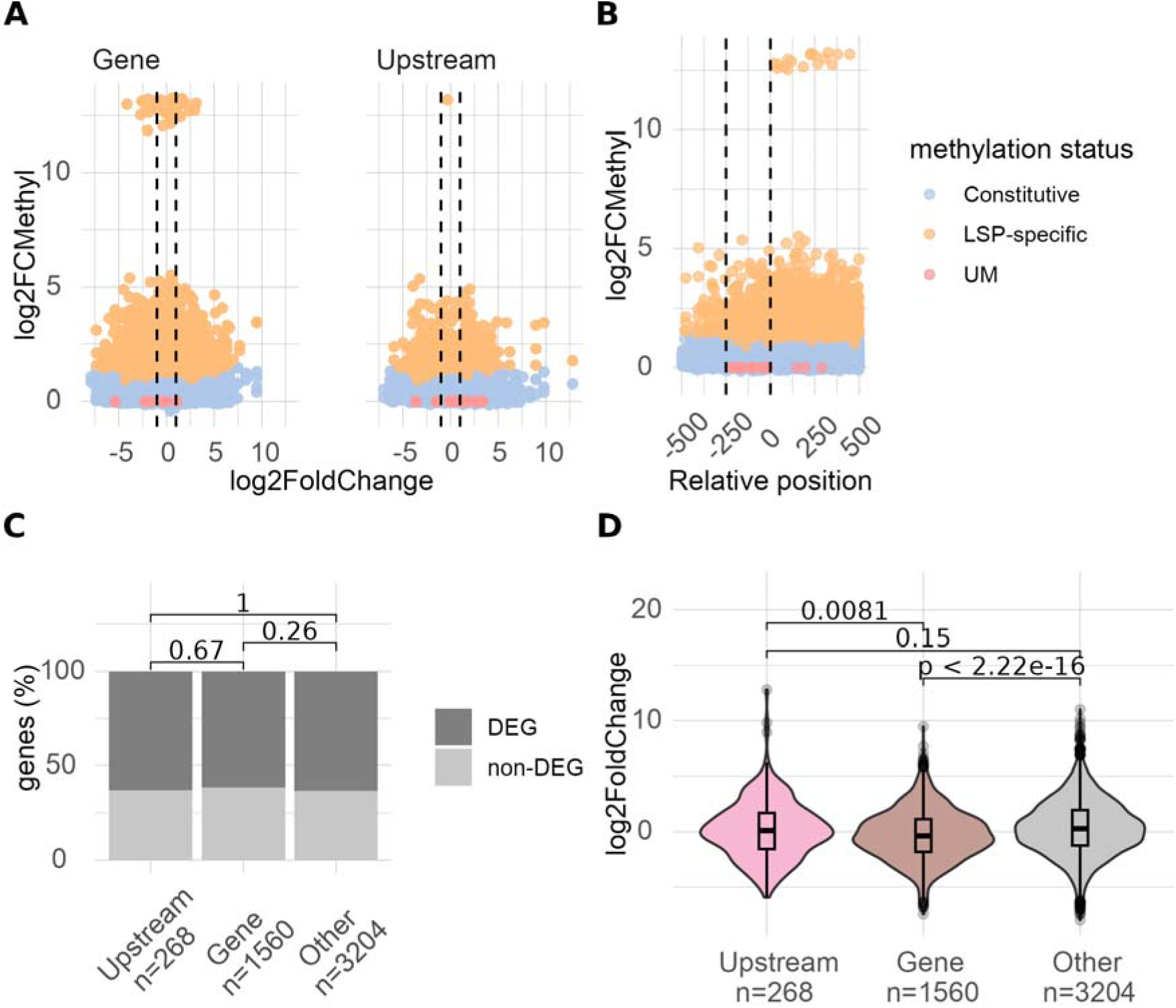
Associations between DNA methylation and gene expression changes between MEP and LSP phases for CCWGG motif. **A.** Comparison of changes in gene expression (log_2_ fold change) and changes in DNA methylation (log_2_FCMethyl) between LSP and MEP for sites localised in intragenic regions (left panel) and upstream regions (right panel). The thresholds for differentially expressed genes are shown as dashed lines (log_2_ fold change is -1 for downregulated and +1 for upregulated genes, padj <0.05) **B.** Relative position of sites (+1 corresponds to the annotated start codon) and changes in DNA methylation between growth phases. There is no statistically significant enrichment of differentially methylated CCWGG sites in the functionally important 0-250 bp upstream region (chi-square test pvalue=0.17). **C.** Qualitative comparison of differentially expressed genes (DEGs) and non-DEGs among groups defined by the presence of differentially methylated sites in upstream regions, intragenic regions, or neither. Genes with a log_2_ fold change ≥ 1 or ≤ –1 and an adjusted p < 0.05 were classified as DEGs; all others were considered non-DEGs. Statistical significance was assessed using a chi-square test. **D.** Quantitative analysis of log_2_ fold change distributions in gene expression for the same groups. Statistical significance was assessed using the Mann–Whitney–Wilcoxon test.

Genes containing LSP-specific CCWGG sites within their intragenic regions showed a tendency toward downregulation at LSP (p < 2.2e-16; Fig. 6D). This observation is consistent with previous findings, which reported that the effect of Dcm methylation on *rpoS* and ribosomal protein gene expression is linked to changes in intragenic CCWGG methylation rather than upstream sites [13,27]. The observed association of differentially methylated intragenic sites and transcriptional downregulation suggests that Dcm methylation may influence gene expression through a mechanism distinct from modulated transcription factor binding to promoter sequences. Further experiments will be required to determine the underlying mechanism.

Only ∼50% of ATGCAT sites were methylated (**Supplementary Table 4**; **Fig. 5**). We identified 103 MEP-specific and 50 LSP-specific sites. No statistically significant associations were found between the presence of condition-specific ATGCAT sites in upstream regions or intragenic regions and changes in gene expression (**Fig. 7**). Unmethylated ATGCAT sites were more frequently located in intergenic regions compared to constitutively methylated or condition-specific sites (**Fig. 3C**), suggesting that they may be associated with transcriptional regulation under conditions not tested here. Consistent with this, Bourgeois *et al*. reported that ATGCAT sites were frequently differentially methylated under infection-related conditions [15], but these changes did not translate into detectable transcriptomic effects. Taken together, these findings highlight that although ATGCAT methylation is dynamic, the biological role of this methylation remains unclear.

**Fig 7.**
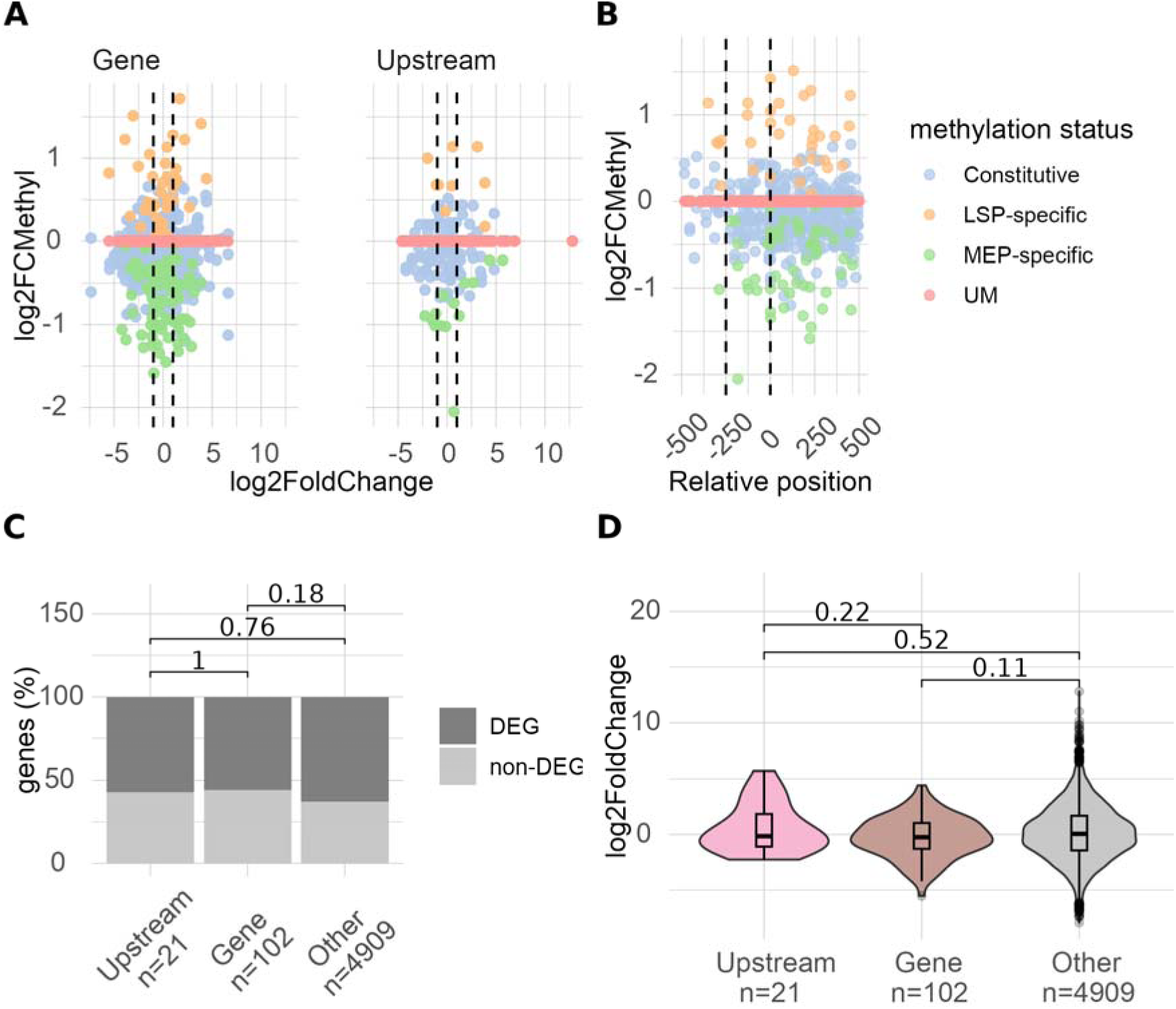
Associations between DNA methylation and gene expression changes between MEP and LSP phases for ATGCAT motif. **A.** Comparison of changes in gene expression (log□ fold change) and changes in DNA methylation (log_2_FCMethyl) between LSP and MEP for sites localised in intragenic regions (left panel) and upstream regions (right panel). The thresholds for differentially expressed genes are shown as dashed lines (log_2_ fold change is -1 for downregulated and +1 for upregulated genes, padj <0.05) **B.** Relative position of sites (+1 corresponds to the annotated start codon) and changes in DNA methylation between growth phases. **C.** Qualitative comparison of differentially expressed genes (DEGs) and non-DEGs among groups defined by the presence of differentially methylated sites in upstream regions, intragenic regions, or neither. Genes with a log_2_ fold change ≥ 1 or ≤ –1 and an adjusted p < 0.05 were classified as DEGs; all others were considered non-DEGs. Statistical significance was assessed using a chi-square test. **D.** Quantitative analysis of log_2_ fold change distributions in gene expression for the same groups. Statistical significance was assessed using the Mann–Whitney–Wilcoxon test.

### 4. Unmethylated sites of regulatory MTases

Unmethylated sites may indicate regions of active regulation and represent potential sources of methylation heterogeneity and epigenetic switching [12]. Comparison of the distributions of constitutively methylated sites and UM sites between intragenic regions and intergenic regions shows statistically significant enrichment of UM sites in the intergenic regions for sites of all regulatory MTases (see **Fig. 4**, **5**, **6**).

For the GATC motif, we identified 44 unmethylated sites on a chromosome (**Supplementary Table 5**), 35 of which were located in intergenic regions. To compare our data with previous methylome studies, we analysed two SMRT sequencing datasets of *S. enterica* ATCC 14028: Sánchez-Romero *et al.* [12] (set1), where growth conditions were not specified, and Bourgeois *et al.* [15] (set2), which analysed cells at the late exponential phase. We identified homologous GATC sites in *S. enterica* ATCC 14028 for 31 out of 35 sites (**Fig. 8C**). In total, 24 of the 31 intergenic unmethylated sites (77%) identified in this study were also reported as unmethylated in at least one of these datasets (**Fig. 8**; **Supplementary Table 9**).

**Fig. 8.**
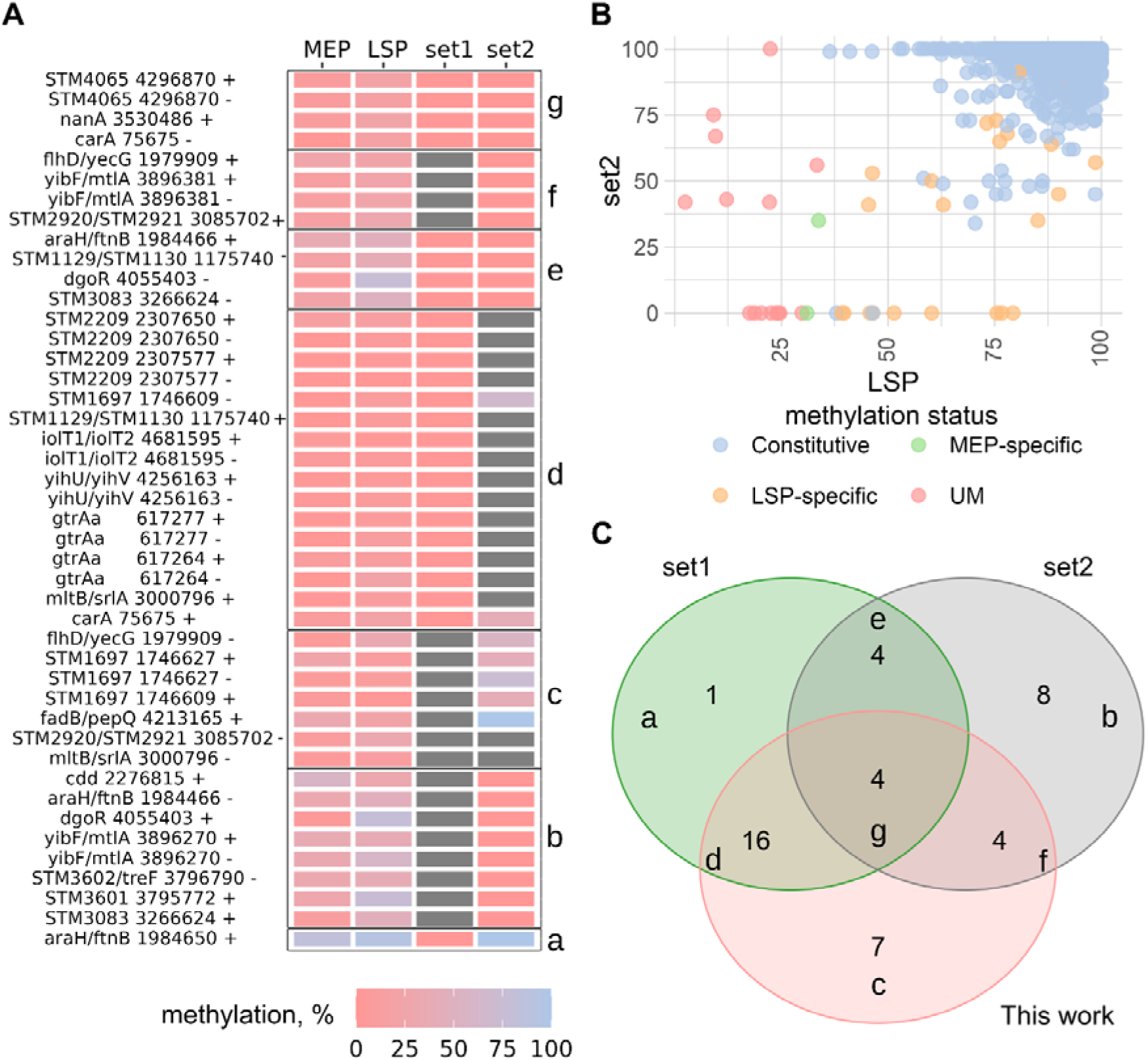
Overlap of unmethylated GATC sites in intergenic regions across three independent datasets. (A) Methylation levels of homologous GATC sites in the chromosomal alignment of *S. enterica* ATCC 14028s and *S. enterica* 4/74, including sites unmethylated in at least one of the three datasets: this study, set 1 [12], and set 2 [15]. Regions labelled a–g correspond to the segments of the Venn diagram shown in panel (C). Grey colour indicates sites for which methylation status was not detected in the corresponding study. (B) Comparison of methylation levels (%) for all homologous GATC sites between *S. enterica* ATCC 14028s in late exponential phase, SMRT data from [15], and our LSP dataset. (C) Venn diagram showing overlap of undermethylated GATC sites among the three datasets.

Four sites were undermethylated in both previous studies but not in ours; three were LSP-specific in our data and one showed low methylation (37–46%), indicating that growth phase largely explains these discrepancies. Among the remaining sites, those unique to Sánchez-Romero *et al.* were fully methylated in our and the Bourgeois *et al.* datasets, while most Bourgeois-specific sites were growth phase–dependent. Only one site showed inconsistent methylation across datasets independent of growth conditions.

No overlap was observed among unmethylated intragenic sites (**Supplementary File 3**). Overall, our Oxford Nanopore data show strong concordance with previous SMRT methylomes and indicate a conserved core of unmethylated GATC sites, whereas sites uniquely unmethylated in one dataset likely reflect lineage-specific Dam methylation patterns or differences in detection sensitivity between technologies.

Unmethylated CCWGG sites are enriched in the upstream regions (Fisher’s exact test pvalue = 2.3e-08). Unmethylated CCWGG are found in the upstream region of ten genes, including *marR, marC, STM1933, STM1934, STM2179, STM2180, yggM, nepI, STM3777, phnS* (see **Supplementary Table 8**). These genes are involved in different stress responses, but the regulatory role of Dcm MTase is unknown.

We identified 162 unmethylated ATGCAT sites in intergenic regions and 462 sites in intragenic regions (**Supplementary Table 7**). Comparison with the dataset from Bourgeois *et al.* [15] (Set 2; **Supplementary File 4**) provided methylation information for 77 of the 162 intergenic sites and 267 of the 462 intragenic sites. Nearly all sites that were unmethylated in our dataset were also unmethylated in the Bourgeois *et al.* dataset, including 97% of intergenic sites and 99% of intragenic sites. In contrast, a substantial proportion of sites in both intergenic and intragenic regions were unmethylated in the Bourgeois *et al.* dataset but not in ours, indicating notable dataset-specific differences in methylation patterns (**Supplementary File 4**). This discrepancy may reflect methodological differences between PacBio and Oxford Nanopore methylation detection, however, we do not observe a similar bias for GATC methylation (**Fig. 8**, **Supplementary File 3**). An alternative explanation is that ATGCAT methylation is inherently less stable and exhibits greater variability between strains. Consistent with this idea, ATGCAT methylation was reported to be the most variable methylation type across conditions in Bourgeois *et al.* [15].

### 5. Methylation heterogeneity

Incomplete DNA methylation generates heterogeneity within a bacterial population by producing subpopulations with distinct methylation states, which can be associated with different phenotypes [12]. This effect is particularly pronounced when multiple methylation sites occur within the same regulatory region, as their combinatorial methylation patterns can create several distinct subpopulations.

A substantial fraction of differentially methylated (32 of 77) and unmethylated (43 of 53) GATC sites are in 36 intergenic regions (**Supplementary Table 9**). 1/36 (2.8%) contained only MEP-specific sites, 3/36 (8.3%) contained only LSP-specific sites, 7/36 (19%) contained only unmethylated sites, and the remaining 25/36 (69.4%) harboured multiple upstream GATC sites with mixed methylation states across conditions (**Supplementary Table 10**). The presence of these GATC clusters upstream of genes suggests potential for epigenetic switches, such as in *E. coli pap* operon [28] or *S. enterica opvAB* [9].

An intergenic region between the *yibF* gene, encoding glutathione S-transferase, and the *mtlA* gene, encoding the mannitol-specific enzyme II of the phosphotransferase system, illustrates this heterogeneity (**Fig. 9**). This region contains three pairs of GATC sites: one classified as constantly methylated, a second as LSP-specific/constantly methylated, and a third as unmethylated in our data. At the LSP, the methylation level of the second GATC site increased at both strands, accompanied by a four-fold increase in *yibF* expression (see **Supplementary Table 1**). Further experiments are required to determine whether methylation of the GATC-2 and GATC-3 sites contributes to the epigenetic regulation of *yibF* and *mtlA*, respectively.

**Fig 9.**
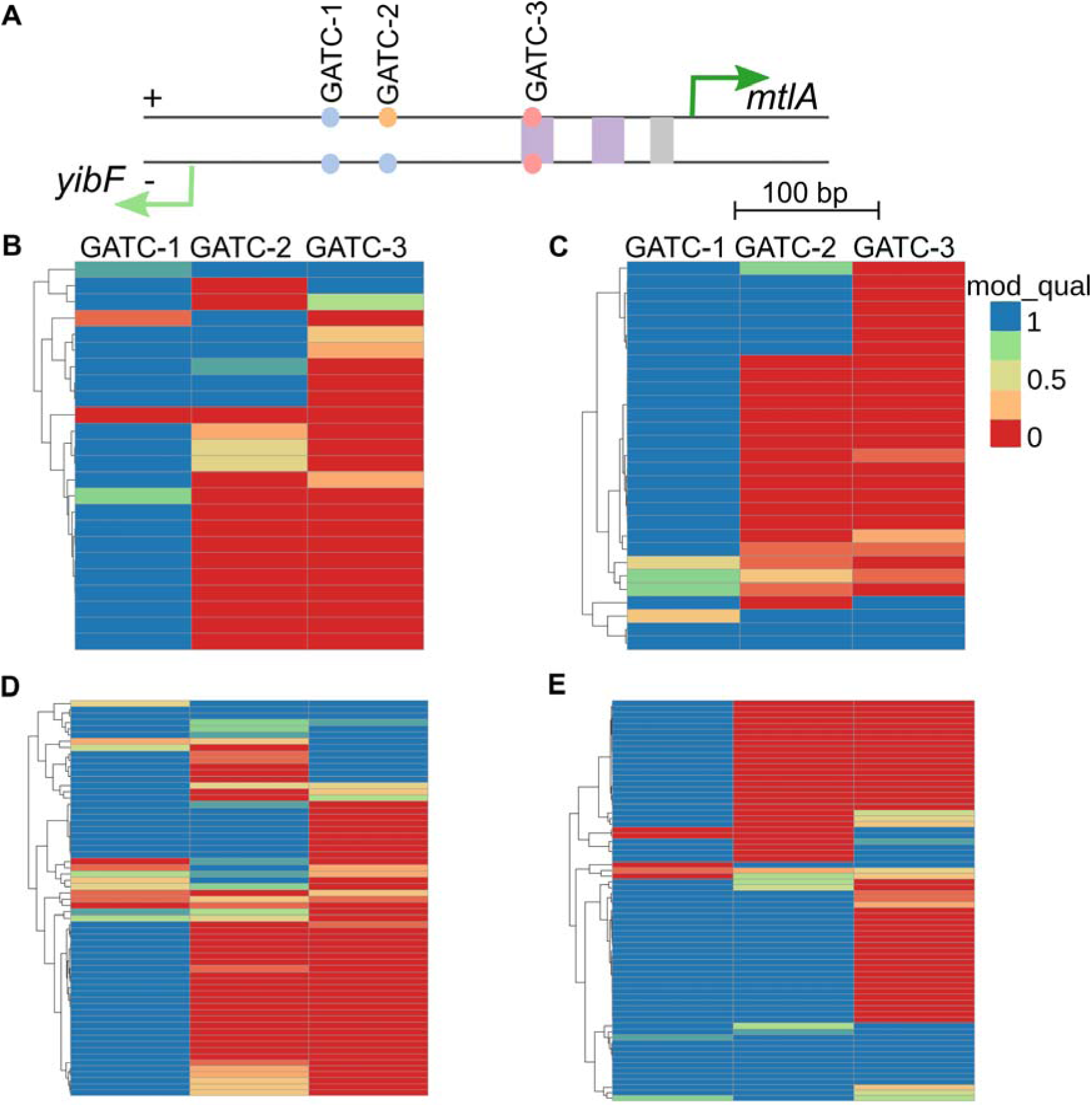
Methylation of GATC sites in the *yibF–mtlA* intergenic region (ST4-74.fa, positions 3,896,108–3,896,582). **A.** Organisation of the intergenic region. Transcription start sites (TSSs) are shown as arrows. GATC sites are depicted as circles, with colors indicating methylation status: blue = constitutively methylated, yellow = LSP-specific methylation, pink = unmethylated. Known transcription factor binding sites are shown as rectangles: violet = Crp binding sites, grey = FruR binding sites according to RegPrecise [29]. **B.** Methylation patterns in the mid-exponential phase (positive strand). **C.** Methylation patterns in the MEP (negative strand). MEP-2 sample, 32× is shown. **D.** Methylation patterns in the LSP (positive strand). **E.** Methylation patterns in the LSP (negative strand). LSP-1, 124× is shown. The heatmap shows the probability of m6A methylation at each GATC site, as determined by *modkit extract-full* (mod_qual column). Each row represents an individual sequencing read.

## Discussion

In this study, we performed a combined analysis of DNA methylation patterns and gene expression changes in *Salmonella enterica* 4/74 between the mid-exponential (MEP) and late stationary (LSP) growth phases. Consistent with previous observations on bacterial epigenomic dynamics [2,30], we found that the regulatory methyltransferases Dam, Dcm, and YhdJ exhibited phase-dependent variation in their methylation patterns, whereas methyltransferases associated with restriction–modification systems maintained consistently high methylation levels at both phases. In addition, we identified sets of completely unmethylated sites for each of the regulatory MTases, which were preferentially located in upstream regions of genes, supporting the idea that methylation can act as a context-dependent regulator of transcription.

From a methodological perspective, our results also demonstrate that Oxford Nanopore sequencing provides a reliable and high-resolution view of the bacterial methylome. The methylation patterns detected using Nanopore were in close agreement with those obtained by PacBio and bisulfite sequencing in previous studies, even at the level of methylation of individual sites. These results are also in line with previous studies reporting on Oxford Nanopore performance to detect methylation in bacteria [17,31].

Most of MTases were relatively lowly expressed at both growth phases but showing further downregulation at LSP compared to MEP. However, we show that their expression does not correlate with their methylation pattern. Perhaps, the methylation pattern is influenced by MTase activity and stability at the protein level rather than by transcription of corresponding genes. Bacteria do not have methylation erasers, and at stationary phase, where cell division is a rare event, even a small amount of active MTases can be enough to methylate the genome. We do not know studies analysing the lifetime of regulatory MTases and their activities, but stability of restriction endonucleases can significantly vary [32], therefore this would be an interesting subject to investigate. A limitation of this study is the difference in sequencing depth between the LSP and MEP samples (124× and 24× vs 32× and 16×, respectively), which may reduce sensitivity for detecting low-coverage methylation sites. Despite this limitation, we observed distinct methylation patterns across all analysed motifs and identified growth-phase-specific differences in methylation that were associated with changes in gene expression. In addition, we recapitulated previously described features, including the enrichment of unmethylated sites in regulatory regions, supporting the biological relevance of the detected methylation patterns.

We analysed the presence of undermethylated and constitutively methylated sites of the *S. enterica* regulatory MTases Dam, Dcm, and YhdJ and assessed their effect on changes in gene expression between MEP and LSP. Binary analysis of differentially expressed genes (DEGs, log_2_FC ≥ 1 or ≤ –1, padj<0.005) and differentially methylated sites did not reveal enrichment in any of the gene groups classified by methylation status (undermethylated sites, constitutively methylated sites, or all remaining genes; **Fig. 9A**, **10A**, **11A**). However, for the GATC motif methylated by Dam MTase, we observed that the genes with LSP-specific methylated sites in their upstream regions were upregulated at LSP compared to the remaining genes. In contrast, genes with LSP-specific sites with a CCWGG motif methylated by Dcm MTase in their intragenic regions were downregulated at LSP compared to the remaining genes. Differentially methylated ATGCAT sites (YhdJ MTase) were not associated with changes in gene expression of the corresponding genes.

Together, these results indicate that DNA methylation can influence the expression of a subset of genes, although its overall impact is subtle. Our findings highlight the value of integrating methylomic and transcriptomic analyses to identify gene subsets that may be subject to epigenetic regulation under specific conditions. These results are in line with recent findings on Dam-dependent epigenetic regulation of *S. enterica* ATCC 14028S genes involved in the oxidative stress response [33], where Zhang *et al.* also analysed changes in methylomic and transcriptomic data. Yang *et al.* found that Dam methylation is dynamic during bacteria cell adaptation to intracellular conditions [25]. While concurrent changes in methylation and gene expression do not necessarily imply a direct regulatory relationship [12,34], condition-specific methylation shifts at genomic sites can still provide important insights into the dynamic regulatory landscape of bacterial genomes.

The identified relatively subtle effect of methylation on gene expression may reflect the existence of differentially methylated subpopulations within the bacterial culture. Under different conditions, the relative abundance of these subpopulations might shift, but rarely to the extent of reaching zero. Consequently, the overall (integrated) gene expression measured across the population differs only slightly between conditions, potentially masking stronger effects at the subpopulation level. Only single-cell transcriptomic and methylomic analyses of bacterial cells in different conditions may identify if there are any differences in gene expression at subpopulation level, however, this is technically not yet feasible.

## Conclusions

Oxford Nanopore data are comparable to other methylation identification methods even at the level of individual sites. DNA methylation in *Salmonella* is dynamic during growth phases and there are associations between changes in DNA methylation and changes in gene expression for Dam and Dcm MTases. These associations demonstrate differences in potential mechanisms of epigenetic regulation, Dam methylation is associated with changes in gene expression if the differentially methylated GATC site is localised in the upstream regions, and Dcm methylation is associated with changes in gene expression if differentially methylated site is localised in the intragenic regions.

## Materials and Methods

### Bacterial strains and growth conditions

*Salmonella enterica* serovar Typhimurium 4/74 [35,36] were maintained on lysogeny broth (Lennox, L-) agar plates (10 g/l tryptone, 5 g/l yeast extract, 5 g/l NaCl, 15 g/l agar). Over-night cultures (16 h) grown in tubes were diluted (1:100) in fresh 25 mL of LB media in 250 mL flasks up to the required growth stages (MEP corresponds to OD_600_ 0.3 and LSP is 16 hours of growth, 220 rpm, 37 °C).

### Extraction of genomic DNA

Genomic DNA was isolated using a phenol:chloroform extraction protocol adapted from [37] . Briefly, 1 mL of overnight culture or 3 mL of OD_600_ 0.3 was pelleted and resuspended in TE buffer (pH 8.0) with 1 mg/mL lysozyme, followed by incubation at room temperature for 5 min. Cell lysis was achieved by adding SDS (final concentration 1.25%) and proteinase K (1 mg/mL), followed by incubation at 55 °C for 2 h. DNA was extracted by phenol:chloroform (1:1) and a chloroform extraction. DNA was precipitated with 3 M sodium acetate (pH 5.2) and isopropanol, washed twice with 70% ethanol, and resuspended in TE buffer (pH 8.0). Residual RNA was removed by treatment with RNase A (Thermo Scientific, EN0531) at 37 °C for 30 min. DNA was cleaned up with an additional phenol:chloroform extraction and ethanol precipitation. The final DNA pellet was resuspended in nuclease-free water. DNA quantity and purity were assessed using a NanoDrop spectrophotometer and Qubit fluorometer (Thermo Fisher Scientific, USA).

### *In vitro* M.EcoRI methylation

1U of M.EcoRI (NEB, #M0211S) was used according to the manufacturer instructions for four hours at 37 °C. DNA was cleaned up with an additional phenol:chloroform extraction and ethanol precipitation.

### Nanopore sequencing

Nanopore sequencing libraries were prepared according to the genomic DNA Ligation Sequencing Kit V14 (SQK-NBD114-24) protocol (Oxford Nanopore Technologies, Oxford Science Park, OX4 4DQ, UK). Prepared libraries were loaded on MinION flow cells (R10.4) and sequenced with the MinION Mk1B device using Kit 14 chemistry and MinKNOW v23.07.8 (Oxford Nanopore Technologies, Oxford Science Park, OX4 4DQ, UK).

### Base calling of nanopore reads

Basecalling of raw Oxford Nanopore signal data was performed using Dorado v0.8.3 with the super accuracy (sup) model dna_r10.4.1_e8.2_400bps_sup@v5.0.0 (5 kHz). Methylation calling for N6-methyladenine (m6A) and N4/N5-methylcytosine (m4C/m5C) was conducted using the corresponding modified base models, dna_r10.4.1_e8.2_400bps_sup@v5.0.0_6mA@v2 and dna_r10.4.1_e8.2_400bps_sup@v5.0.0_4mC_5mC@v2. Demultiplexing was performed using the dorado demux command with the kit name specified as --kit-name SQK-NBD114-24. Bam files were aligned on a *Salmonella enterica* 4/74 reference genome (GenBank IDs: CP002487.1 - CP002490.1) using Dorado aligner and sorted by Samtools v1.18. Sorted ModBAM files were analyzed using modkit v0.4.4. Methylated positions in a genome were identified using modkit pileup with automatic threshold detection based on the input data, and motifs were identified using modkit find-motifs with default parameters. Motif occurrences were quantified separately for each DNA strand, allowing strand-specific analysis of methylation patterns. modkit extract full was used to extract information on the base modification probabilities.

### DNA methyltransferases and R-M systems

We used REBASE annotation and PacBio data to identify R-M systems in the 4/74 genome and their methylation sites.

### Analysis of site distribution across the genome

Seqkit locate with the --bed option (v2.1.0) [38] was used to identify the genomic coordinates of sites corresponding to each methylated motif. To analyse motif methylation, the resulting coordinates were intersected with bedMethyl files using bedtools intersect (v2.23.0) [39] with the -s option.

### Site annotation

Methylated sites were annotated using the *S. enterica* 4/74 annotation from [21], which includes sRNAs (**Supplementary File 5**). Using genome annotation in GFF format, genomic coordinates were classified as either intragenic regions (including coding sequences and non-coding RNAs) or intergenic regions. Chromosomal macrodomains were annotated based on the definitions provided in Cameron *et al*. [40]. If a macrodomain boundary fell within a gene, the macrodomain was extended to the end of that gene (**Supplementary Table 6**). Sites in intragenic regions assigned to the corresponding genes. Sites in intergenic regions are assigned to the downstream genes as “upstream” sites and to the downstream genes as “downstream” sites. The analysis of site distribution patterns was performed by custom R script.

### Methylation threshold identification

To establish an optimal threshold for calling a site methylated based on Oxford Nanopore signal, we performed a receiver operating characteristic (ROC) analysis using the GAATTC (m6A) motif targeted by M.EcoRI as a ground-truth reference. Genomic DNA from *Salmonella enterica* serovar Typhimurium 4/74 (Late Stationary Phase) was sequenced both before and after in vitro methylation with M.EcoRI, which fully methylates GAATTC sites. These paired datasets provided unmethylated (untreated) and fully methylated (treated) controls for threshold calibration. Methylation levels of all GAATTC sites were evaluated at candidate thresholds from 0% to 100% in 10% increments. For each threshold, sites were classified as methylated or unmethylated. We calculated True Positives (TP) as sites called methylated at the given threshold and methylated in the M.EcoRI-treated sample; True Negatives (TN) as sites called unmethylated and unmethylated in the untreated sample; False Positives (FP) as sites called methylated in the untreated sample; and False Negatives (FN) as sites called unmethylated in the treated sample. Sensitivity was calculated as TP / (TP + FN), and specificity as TN / (TN + FP). ROC analysis was performed using the pROC package (version 1.18.5) in R. The analysis identified a 35% methylation level as the optimal classification threshold, yielding specificity = 0.992 and sensitivity = 0.996. This threshold was applied in all subsequent analyses of methylation state.

### Sequencing depth effect analysis

A sample sequenced at 120× depth was used to extract all GATC sites, which were binned according to their methylation levels. To evaluate the effect of coverage on methylation accuracy, we generated 10 random subsamples for each target depth from 5× to 90×. For every subsample, the methylation level of each site was recalculated, and the deviation from the original 120× estimate was determined. This allowed us to assess how reduced sequencing depth influences the stability and accuracy of methylation quantification.

### Transcriptomic data analysis

Transcriptomic data of MEP, ESP, and LSP growth stages was used for wild-type *S. enterica* 4/74 published earlier [21]. Bowtie2 (2.5.2) was used to map the raw fastq files on *S. enterica* ST4/74 genome [41]. featureCounts was used to get the count table. *S. enterica* 4/74 annotation from Kröger *et al*. [21], which includes sRNAs (**Supplementary File 5**) was used as an annotation file. Differentially expressed genes for growth stage data (ESP vs MEP and LSP vs MEP, two biological replicas for all stages) were identified using the DESeq2 package v1.49.3. TPM values were calculated by custom R script. Three growth stages (MEP, ESP, and LSP) were used for MTase and R-M system gene expression analysis. Two growth stages, MEP and LSP, were used for integrative methylomic and transcriptomic analysis.

### Comparison with data on *S. enterica* ATCC 14028

Data on undermethylated GATC sites [12] and on GATC and ATGCAT methylation [15] were used for comparison. Both studies were performed on *S. enterica* ATCC 14028. Chromosomes of ATCC 14028 (GenBank ID: CP001363) and 4/74 were aligned using progressive Mauve, core genome blocks were aligned with MAFFT v7.505, and homologous site coordinates were identified with an in-house Python script. Methylation levels at homologous sites were then compared between the two genomes.

### Statistical analysis

The Fisher’s exact test/Chi-square test was used for categorical data. Wilcoxon–Mann–Whitney tests were performed when comparing distributions.

## Data availability

The data have been deposited with links to BioProject accession number PRJNA1380450 in the NCBI BioProject database (https://www.ncbi.nlm.nih.gov/bioproject/). The processed data are available at https://github.com/asershova/salmonella_methylation_paper_2025.

## Code availability

The data analysis in this study was performed with R and Python code, available at https://github.com/asershova/salmonella_methylation_paper_2025.

## Acknowledgements

Deirdre Muldowney (TCD) is acknowledged for technical support.

## Author contributions

Conceptualisation (ASE, CK), data curation (ASE), formal analysis (ASE), investigation (ASE, CH, CK), methodology (ASE), resources (ADSC, KH), supervision (CK), writing original draft (ASE, CK), writing – review and editing (ASE, CH, KH, ADSC, CK).

## Supporting Information

Supplementary File 1. Methylation detection thresholds and the effect of sequencing depth on methylation detection

Supplementary File 2. Methylated ATGCAT genomic context and R-M system MTase methylation analysis

Supplementary File 3. Comparative analysis of GATC methylation in intragenic regions

Supplementary File 4. Comparative analysis of ATGCAT methylation

Supplementary File 5. Genome annotation of *S. enterica* ST4-74 used in this work

Supplementary Table 1. DESeq2 data (LSP vs MEP)

Supplementary Table 2. modkit identified motifs

Supplementary Table 3. modkit identified motif distribution across samples

Supplementary Table 4. Motif methylation level

Supplementary Table 5. Whole genome GATC site methylation

Supplementary Table 6. Site distribution by macrodomains

Supplementary Table 7. Whole genome ATGCAT site methylation

Supplementary Table 8. Whole genome CCWGG site methylation

Supplementary Table 9. Intergenic regions containing unmethylated and conditionally methylated GATC sites, pivot table

Supplementary Table 10. Intergenic regions containing unmethylated and conditionally methylated GATC sites

