## Supplementary material for "Methylome and transcriptome mapping reveal miniscule DNA methyltransferase regulons in *Salmonella enterica* serovar Typhimurium": Suppl.File 1

### **Supplementary File 1.**

### **1. Evaluation of Nanopore-Based Methylation Detection Using *in vitro* methylation**

To evaluate the sensitivity and accuracy of Nanopore-based methylation detection, we compared genomic DNA from *S. enterica* 4/74 *in vitro* treated with M.EcoRI to an untreated control. M.EcoRI specifically methylates the second adenine of the GAATTC recognition sequence. Strain 4/74 does not encode any native methyltransferases (MTases) capable of modifying this motif. Both treated and untreated DNA samples were sequenced, and methylation levels for all nucleotides within GAATTC motif occurrences were quantified using the modkit tool. As expected, the second adenine exhibited the highest methylation signal among all positions within the motif (**Fig. S1.1A**), consistent with the established activity of M.EcoRI. This result demonstrates that Nanopore sequencing combined with modkit analysis reliably detects site-specific DNA methylation and supports the use of M.EcoRI-modified DNA as a positive control for defining a detection threshold.

The first adenine within the GAATTC motif also exhibited elevated methylation levels in the M.EcoRI-treated sample compared to the untreated control. This phenomenon may reflect the known influence of methylating adjacent nucleotides, as reported previously [1]. Nonetheless, methylation at the second adenine—corresponding to the known M.EcoRI target—was markedly higher, confirming the specificity of the modification. However, methylation of the second adenine did not reach 100% in most cases, with a median value of approximately 75% (**Fig. S1.2**). This incomplete methylation may result from suboptimal *in vitro* methylation, influence of local sequence context or secondary structure during signal detection and base calling.

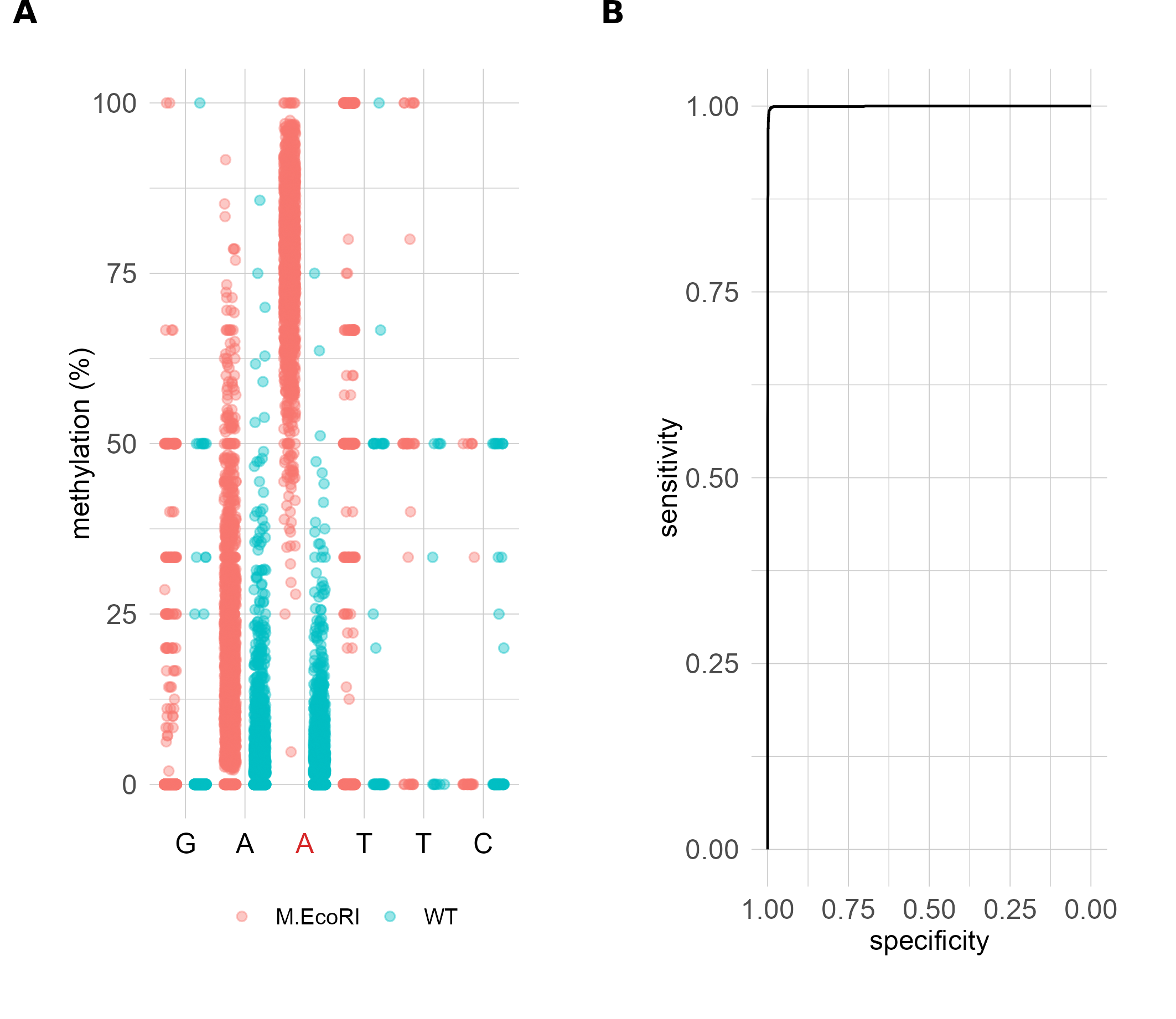

**Fig S1.1. Comparison of M.EcoRI treated and non-treated (WT) samples of *S. enterica* 4/74 genomic DNA.** **A**. Methylation % of bases of GAATTC sites in M.EcoRI treated (M.EcoRI) and non-treated (WT) samples **B**. ROC curve based on the comparison of the methylation level of the 2nd A in the GAATTC sequence between M.EcoRI treated and non-treated samples. Methylation percentage represents the proportion of sequencing reads containing a methylated base at a given genomic position relative to the total number of reads covering that position with the corresponding base.

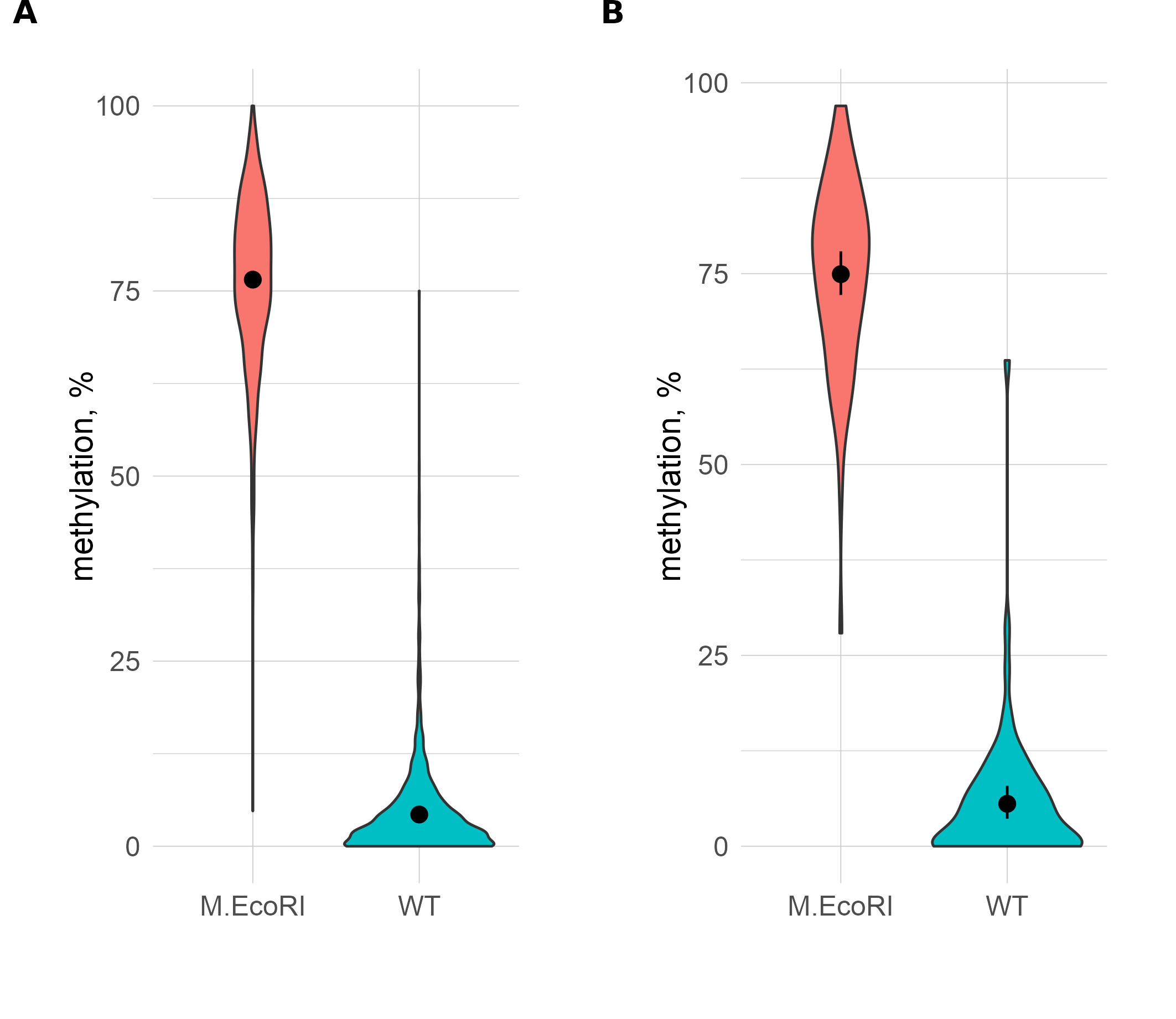

**Fig. S1.2 Methylation of the 2nd A of GAATTC sites of the chromosome (A) and plasmids (B).** Methylation % of the second A of GAATTC sites in M.EcoRI treated (M.EcoRI) and non-treated (WT) samples.

In untreated controls, both adenines within the GAATTC motif exhibited low and comparable methylation levels yet none were 0% (**Fig. S1.1A**). To define a threshold for distinguishing methylated from non-methylated positions, we constructed a receiver operating characteristic (ROC) curve using the second adenine in the GAATTC motif (**Fig. S.1.1B**). This analysis determined that a methylation level threshold of 35% achieves high performance, yielding a specificity of 0.992 and a sensitivity of 0.996.

### 2. **Effect of Sequencing Depth on Methylation Detection**

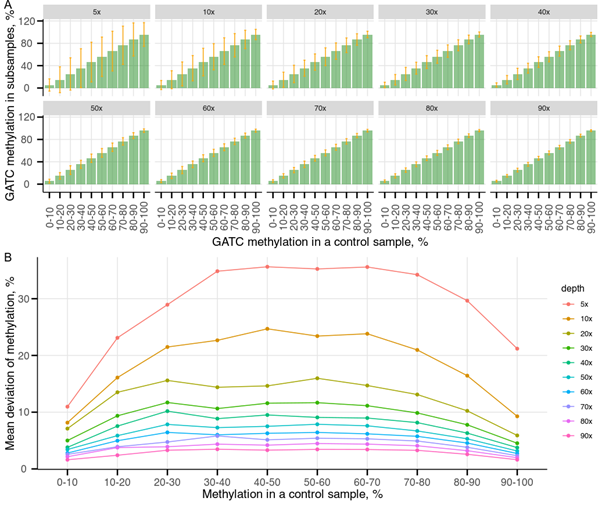

**Fig S1.3. The effect of sequencing depth on accuracy of methylation level estimates across GATC sites in *Salmonella*.** A sample sequenced at 120× depth by samtools was used to extract all GATC sites, which were grouped into bins based on their methylation levels. For each sequencing depth (5× to 90×), 10 random subsamples were generated. The observed methylation level and its deviation from the original value (120× sample) were calculated for each site. **A.** Bar chart showing the average observed methylation level for each bin across sequencing depths. Vertical orange lines represent the mean deviation from the control sample. **B.** Mean deviation from the original methylation level for each bin depending on sample sequencing depth.

GATC sites are methylated by Dam MTase in *S. enterica* and other *Enterobacteriaceae*, with PacBio data from the REBASE database indicating over 99% methylation of these sites in *S. enterica* genomes. To assess how sequencing depth affects methylation detection, we used GATC sites in *Salmonella* as a model. A sample sequenced at 120× depth served as the reference, from which 10 random subsampled datasets were generated for each target depth between 5× and 90× (Fig. 2). Methylation levels at GATC sites were estimated for each depth and compared to the high-depth reference.

Our results indicate that sequencing depth has a differential impact on the resolution of methylation levels. Sites with extreme methylation values (0–20% and 80–100%) remained distinguishable even at low coverage, whereas intermediate levels required substantially higher sequencing depth for reliable separation (Fig. 2A, B). The mean deviation from the reference methylation levels decreased sharply between 5× and 30×, after which further increases in sequencing depth resulted in only modest improvements of methylation estimation (Fig 3). These results are similar to the previous estimations (doi: 10.1101/2024.11.09.622763v1).

*modkit* applies internal quality thresholds for classifying reads as methylated or non-methylated. As a result, not all reads present in the BAM files are included in the methylation analysis performed by *modkit*. **Table S1** shows the mean sequencing depth calculated using *samtools* alongside the mean number of reads utilized by *modkit* for calculating the methylation percentage of the GATC motif. To take into account the differences in sequencing depth of biological replicas, we calculated methylation level using a weighted average, where sequencing depth of the sample was taken into account.

**Table S1**. Comparison of mean sequencing depth (samtools) and mean number of reads used by modkit for methylation calculation at GATC sites on chromosome.

| sample | Samtools meandepth | total reads for A in GATC position in modkit bed files |
| --- | --- | --- |
| MEP-1 | 16.7 | 7.7 |
| MEP-2 | 32.4 | 14.7 |
| LSP-1 | 124.3 | 56.9 |
| LSP-2 | 24.4 | 10 |
