## Supplementary material for "Methylome and transcriptome mapping reveal miniscule DNA methyltransferase regulons in *Salmonella enterica* serovar Typhimurium": Suppl.File 2

1. Methylated ATGCAT context analysis.

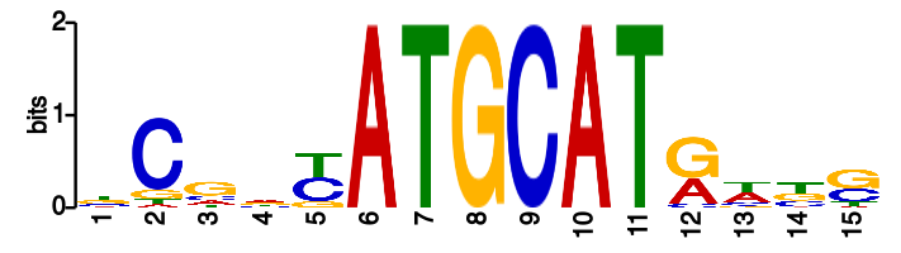

**Fig. S2.1. Methylated ATGCAT context**. Sequences, containing ATGCAT sites and 5 bp from each side with more than 50% of methylation at both phases were used for the motif search by MEME.

According to Fig. 1, there is no specific sequence context, surrounding methylated ATGCAT sites, there is a weak preference of pyrimidines in the position before the site and purins in the position after the site.

2. RM-system MTase methylation

Jitter plots, showing methylation level in different positions of sites (Fig. 7A, C, Fig. 8 A), clearly show the methylated position, and also show that methylation signal can affect neighbouring positions.

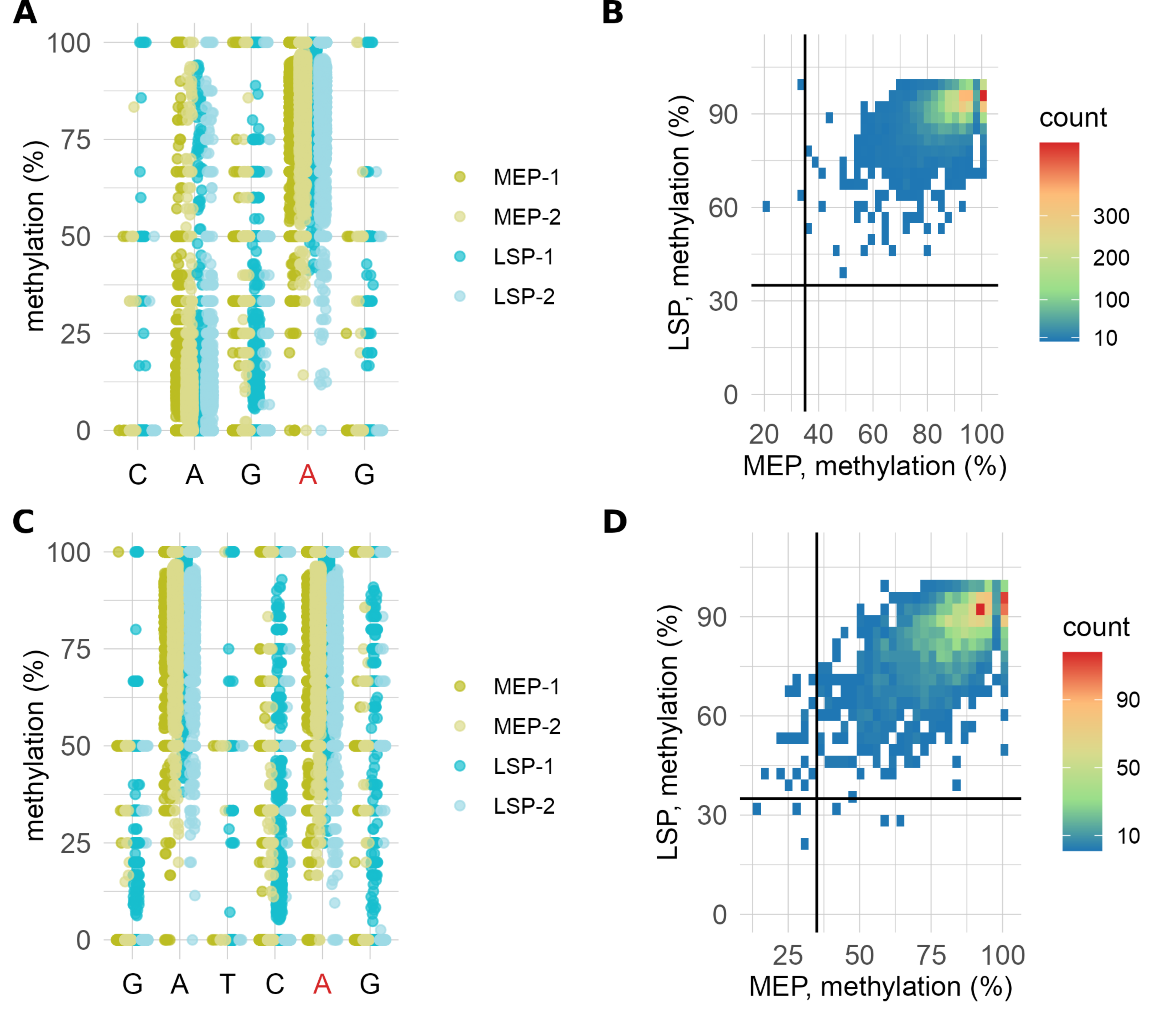

**Fig S2.2. Type III (CAGAG) and Type IIG (GATCAG) R-M methylation. A.** Methylation percentage of bases within CAGAG sites across the genome. The second adenine is predominantly methylated (shown in red). Two Mid-Exponential Phase samples (MEP-1 and MEP-2) and two Late Stationary Phase samples (LSP-1 and LSP-2) are shown. **B.** Methylation dynamics of CAGAG sites (n = 6,125) across growth phases. **C.** Methylation percentage of bases within GATCAG sites across the genome. The second adenine is predominantly methylated (shown in red). The first adenine is methylated because GATCAG contains a GATC motif recognised by the Dam MTase. Two Mid-Exponential Phase samples (MEP-1 and MEP-2) and two Late Stationary Phase samples (LSP-1 and LSP-2) are shown. **D.** Methylation dynamics of GATCAG sites (n = 2,978) across growth phases. Solid lines indicate the 35% methylation threshold.

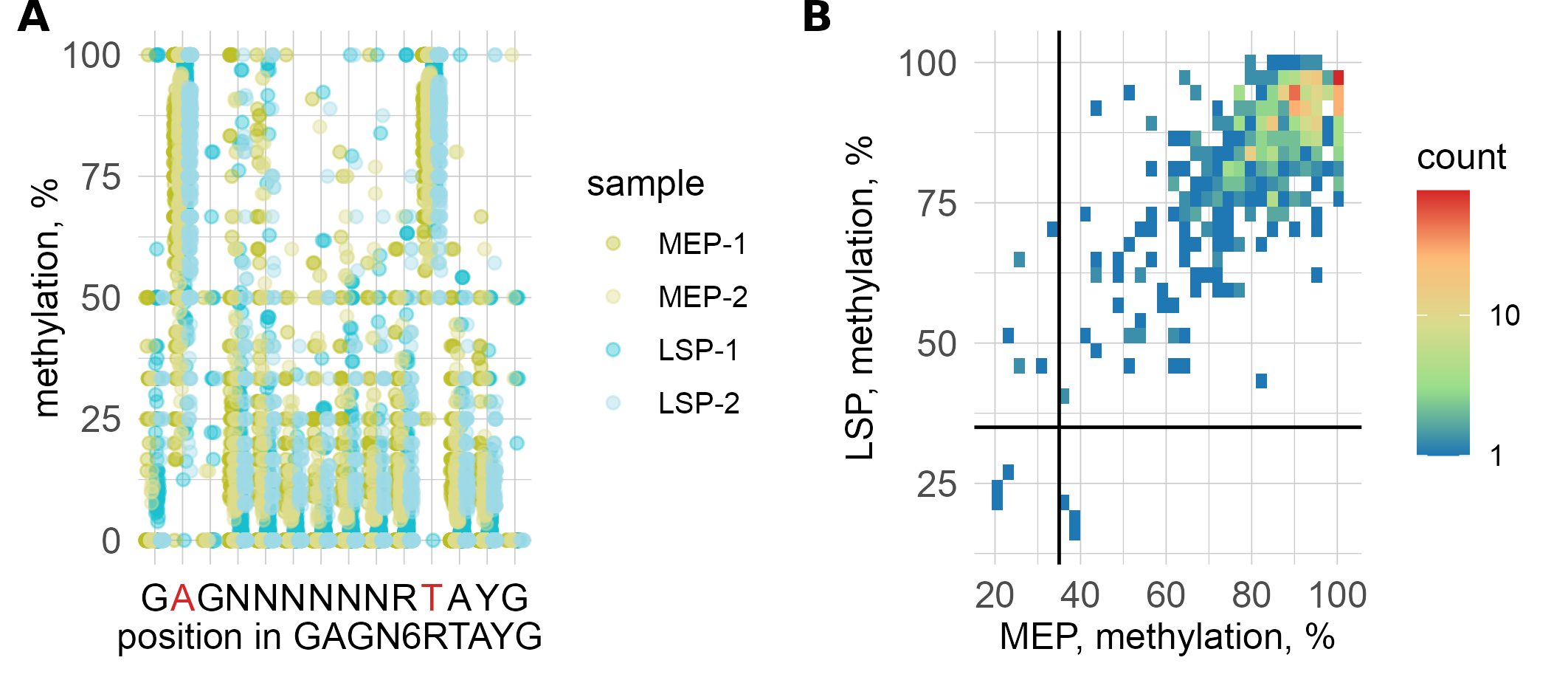
**Fig S2.3. Type I R-M system (GAG(N)_6_RTAYG) methylation. A.** Methylation % of bases in the GAG(N)_6_RTAYG sites. An A in the first position on the direct strand and an A in the 10^th^ position on the reverse strand are methylated (shown in red). Two Mid-Exponential Phase samples (MEP-1 and MEP-2) and two Late Stationary Phase samples (LSP-1 and LSP-2) are shown. **B.** Methylation dynamics of GAG(N)_6_RTAYG sites (n = 504) across growth phases. Solid lines indicate the 35% methylation threshold.
