## Supplementary material for "Methylome and transcriptome mapping reveal miniscule DNA methyltransferase regulons in *Salmonella enterica* serovar Typhimurium": Suppl.File 3

**Supplementary File 3. Comparative analysis of GATC methylation in intragenic regions for three datasets**

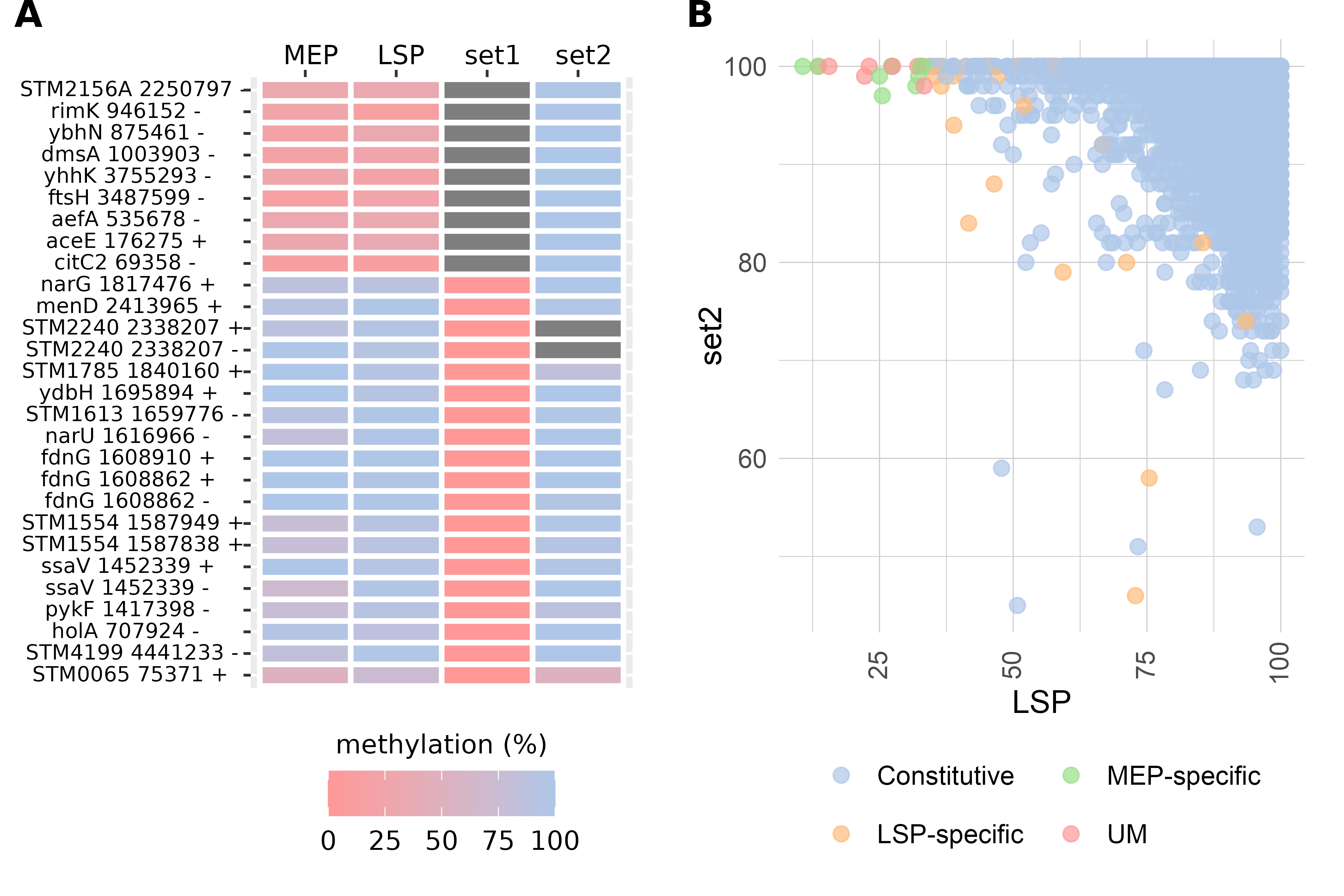

**Fig. S3.1. Overlap of unmethylated GATC sites in intragenic regions across three independent datasets**. (A) Methylation levels of homologous GATC sites in the chromosomal alignment of *S. enterica* ATCC 14028s and *S. enterica* 4/74, including sites unmethylated in at least one of the three datasets: this study, set 1 [PMC7708049], and set 2 [PMC9239280]. (B) Comparison of methylation levels (%) for all homologous GATC sites between *S. enterica* ATCC 14028s in late exponential phase, SMRT data from [PMC9239280], and our LSP dataset.

We identified nine unmethylated intragenic GATC sites in our dataset (**Fig. 3.1A**). In comparison, Sánchez-Romero et al. [PMC7708049] (set 1) reported 19 unmethylated intragenic GATC sites, whereas no intragenic unmethylated GATC sites were detected in the Bourgeois et al. dataset [PMC9239280] (set 2). Direct comparison of methylation levels between our LSP-2 sample and the Bourgeois et al. dataset (**Fig. 3.1B**) showed that most intragenic GATC sites were highly methylated in both datasets. However, several sites displayed moderately lower methylation in our data but higher in set 2, and vice versa. These differences likely reflect a combination of strain-specific variation, growth-phase differences, and the distinct sensitivities of the Oxford Nanopore and SMRT sequencing technologies.
