## Supplementary material for "Methylome and transcriptome mapping reveal miniscule DNA methyltransferase regulons in *Salmonella enterica* serovar Typhimurium": Suppl.File 4

**Supplementary File 4**. **Comparative analysis of ATGCAT methylation in two datasets**

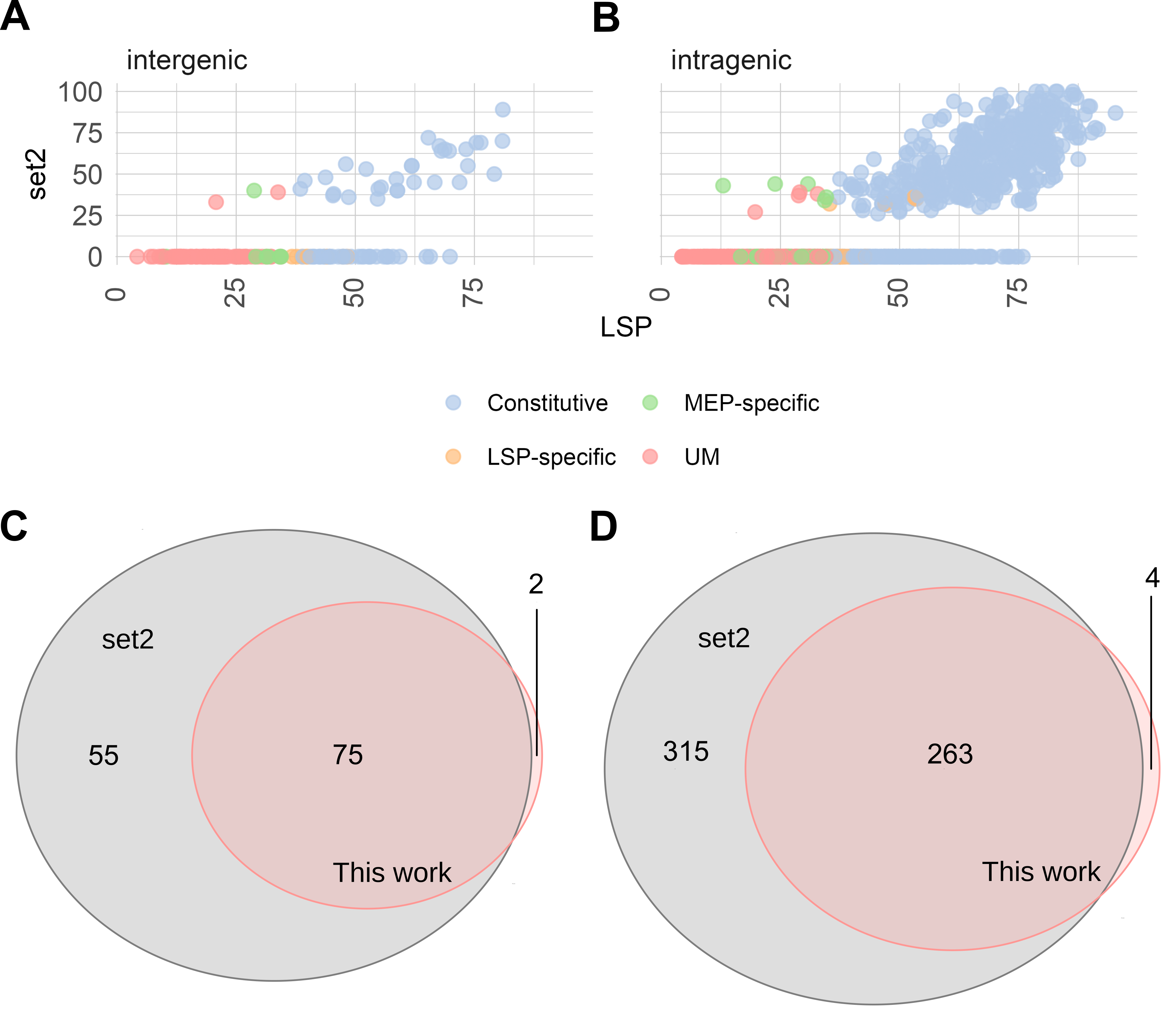

**Fig. S4.1. Comparison of ATGCAT site methylation between our dataset and set 2 [PMC9239280].**
 (A–B) Methylation levels (%) for 1,206 homologous ATGCAT sites in *S. enterica* ATCC 14028s at the late exponential phase, comparing SMRT sequencing data [PMC9239280] with our LSP dataset. Methylation levels are shown separately for intergenic regions (n=170) (A) and intragenic regions (n=1036) (B). (C–D) Overlap of unmethylated ATGCAT sites identified in our dataset and in set 2, shown for intergenic regions (C) and intragenic regions (D).
